# Spectrin coordinates cell shape and signaling essential for epidermal differentiation

**DOI:** 10.1101/2024.10.17.618796

**Authors:** Arad Soffer, Aishwarya Bhosale, Roohallah Ghodrat, Marc Peskoller, Takeshi Matsui, Carien M. Niessen, Chen Luxenburg, Matthias Rübsam

## Abstract

Cell shape and fate are tightly linked, yet how the cortical cytoskeletal integrates regulation of shape and fate remains unclear. Using the epidermis as a paradigm where cell shape changes guide differentiation we identify spectrin as an essential organizer of the keratinocyte actomyosin cortex that integrates transitions in cell shape with spatial organization of signaling. Loss of αII-spectrin (*Sptan1*) in the mouse epidermis altered cell shape in all layers and impaired differentiation, and barrier formation. High-resolution imaging and laser ablation revealed that E-cadherin guides differential gradients of actin and spectrin to regulate the layer-specific organization and mechanics of sub-membraneous spectrin-actomyosin networks. This organization is essential to dissipate tension, maintain structural integrity, and retain active growth factor receptor EGFR and the cation channel TRPV3 at the membrane in upper layers to induce terminal differentiation. Together, these findings reveal how organization of the cortical cytoskeleton coordinates changes in cell shape and cell fate at the tissue scale necessary to establish epithelial barriers.

## Introduction

Cells in our body exhibit a wide range of shapes, sizes and functions, from giant, multinucleated osteoclasts and contractile skeletal muscle cells to small, disc-shaped red blood cells (Luxenburg et al., 2007; Calderón et al., 2014; Elgsaeter et al., 1986). Defects in cell shape often correlate with tissue malfunction and disease phenotypes (Lee et al., 2016; Kalluri and Weinberg, 2009; Delaporte et al., 1990; Emery, 2002; PAULING and ITANO, 1949). For example, in common skin diseases such as psoriasis, atopic dermatitis as well as skin carcinoma, keratinocytes exhibit abnormal cell shape associated with impaired differentiation and compromised epidermal barrier function (Koegel et al., 2009; Van Der Kammen et al., 2017). However, whether cell shape directly regulates cell differentiation and tissue function remains an open question.

At the body’s surface, the interfollicular epidermis (hereafter, epidermis) is a stratified, multilayered epithelium that provides a barrier to protect organisms from water loss and external challenges. The epidermis exhibits progressive cell shape changes that are linked to the differentiation status of keratinocytes. Cuboidal basal stem cells that initiate differentiation and move upwards into the spinous layer increase in size and start to flatten. Further, upon entering the first of the three granular layers (SG3-1 basal to apical), keratinocytes further flatten substantially to then take on the shape of kelvin’s flattened tetrakaidecahedrons in SG2 (Yokouchi et al., 2016) essential to maintain the tight junctional (TJ) barrier in this layer (Kubo et al., 2009; Yoshida et al., 2013; Yokouchi et al., 2016; Rübsam et al., 2017). Terminal differentiation and final flattening coincides with terminal differentiation in the SG1, a process that involves enucleation and transglutaminase (TGM)-mediated crosslinking of structural proteins and barrier lipids resulting in the formation of the dead stratum corneum barrier that physically separates the organism from the environment (Simpson et al., 2011a; Fuchs, 2007). The mammalian epidermis is thus an ideal model to address the relationship between cell position, shape and differentiation.

Cell shape is mainly determined by the actomyosin cytoskeleton and its associated adhesion receptors. The actomyosin cytoskeleton is a contractile subcellular network made of actin fibers (F-actin), myosin II motor proteins, and dozens of actin-binding proteins that regulate its organization, dynamics, and contractile state and thus the mechanical properties (Pollard, 2016). By engaging integrin-based cell– matrix adhesions and cadherin-based adherens junctions (AJs), actomyosin-generated forces are transmitted across cells and tissues (Noordstra et al., 2023; Saraswathibhatla et al., 2023). Moreover, actomyosin activity also modifies signaling cascades involved in fate regulation (Luxenburg and Zaidel-Bar, 2019; Pollard, 2016; Zaidel-Bar et al., 2015). In the epidermis cortical F-actin levels increase with differentiation and cell flattening, a process regulated by E-cadherin (Rübsam et al., 2017). Yet, how cortical actomyosin networks are spatially patterned to drive cell-shape transitions, and how these changes are linked to differentiation, remain poorly understood.

Spectrins are tetramers consisting of two α- and two β-spectrins. Vertebrates have two α-spectrin (αI and αII) and five β-spectrin (βI-βV) variants encoded by different genes. While αI-spectrin is expressed only in erythrocytes, αII-spectrin (encoded by *Sptan1*) is the only α-spectrin in non-erythrocyte cells that can interact with all five β-spectrins (Bennett and Healy, 2009; Bennett and Lorenzo, 2016; Teliska and Rasband, 2021).

Spectrin tetramers are large, tensile, spring-like complexes that integrate into the cortical actomyosin network in cell types such as erythrocytes, neurons and fibroblasts (Leterrier and Pullarkat, 2022). The shock-absorbing, actin-crosslinking capacity of spectrins in combination with a dynamic interplay with cortical myosin allows spectrin–actin networks to generate and sense local forces to not only regulate cell shape but also their elastic properties (Leterrier and Pullarkat, 2022; Lorenzo et al., 2023; Ghisleni et al., 2020). Moreover, the organization of spectrin-actomyosin networks promotes microdomain formation of transmembrane proteins, e.g. ion channels, in the plasma membrane (Lorenzo et al., 2023). The function of Spectrin in epithelia remains less well defined. As in erythrocytes, spectrin controls cortical actomyosin organization and cell shape in *Drosophila* follicular epithelium and in *C. elegans* epidermis (Praitis et al., 2005; Ng et al., 2016). In mice, *Sptan1* knockout is embryonic lethal, characterized by neural tube-, cardiac-and craniofacial defects and abnormal growth (Stankewich et al., 2011). In cultured keratinocytes spectrin was shown to influence early keratinocyte differentiation (Zhao et al., 2011, 2013; Wu et al., 2015) but whether spectrins direct changes in cell shape and regulate epidermal barrier formation is not known.

In the present study, we found that αII-spectrin levels increased during epidermal differentiation to control actomyosin organization, contractility, and structural integrity. Moreover, E-cadherin-dependent integration of spectrin into the submembraneous cortex spatially coordinates cell shape transitions and the activation of an EGFR/TRPV3 signaling pathway required for terminal differentiation and thus epidermal barrier formation. These observations provide novel insights into how cortical organization, mechanical properties, and cell fate regulation are connected in a physiologically relevant mammalian system.

## Results

### αII-spectrin determines epidermal cell shape

Although it is well established that keratinocytes adopt an increasingly flattened squamous morphology during stratification, their exact 3D shapes and volumes have not been defined. To this end, we used a K14-Cre driven rainbow transgene (Snippert et al., 2010) to fluorescently label single cells in vivo followed by 3D rendering of the cytoplasmic YFP signal to determine cell shape, size and volume (Fig. 1a,b; Suppl. Fig. 1a). This analysis showed that initially cuboidal basal keratinocytes double in size and volume when transiting into the spinous layer as well as when moving into the granular layer while preserving surface to volume ratios during stratification. (Fig. 1a-c; Suppl. Fig. 1b). Only when these cells become fully flattened corneocytes the size to volume ratio is increased due to a further increase in surface area (1.41 fold). Moreover, the size and volume in each layer are remarkably stereotypic also between mice (Fig.1a-c, Supple. Fig.1b). Thus, keratinocytes undergo highly stereotypic cell size, shape and volume changes during stratification.

**Figure 1.**
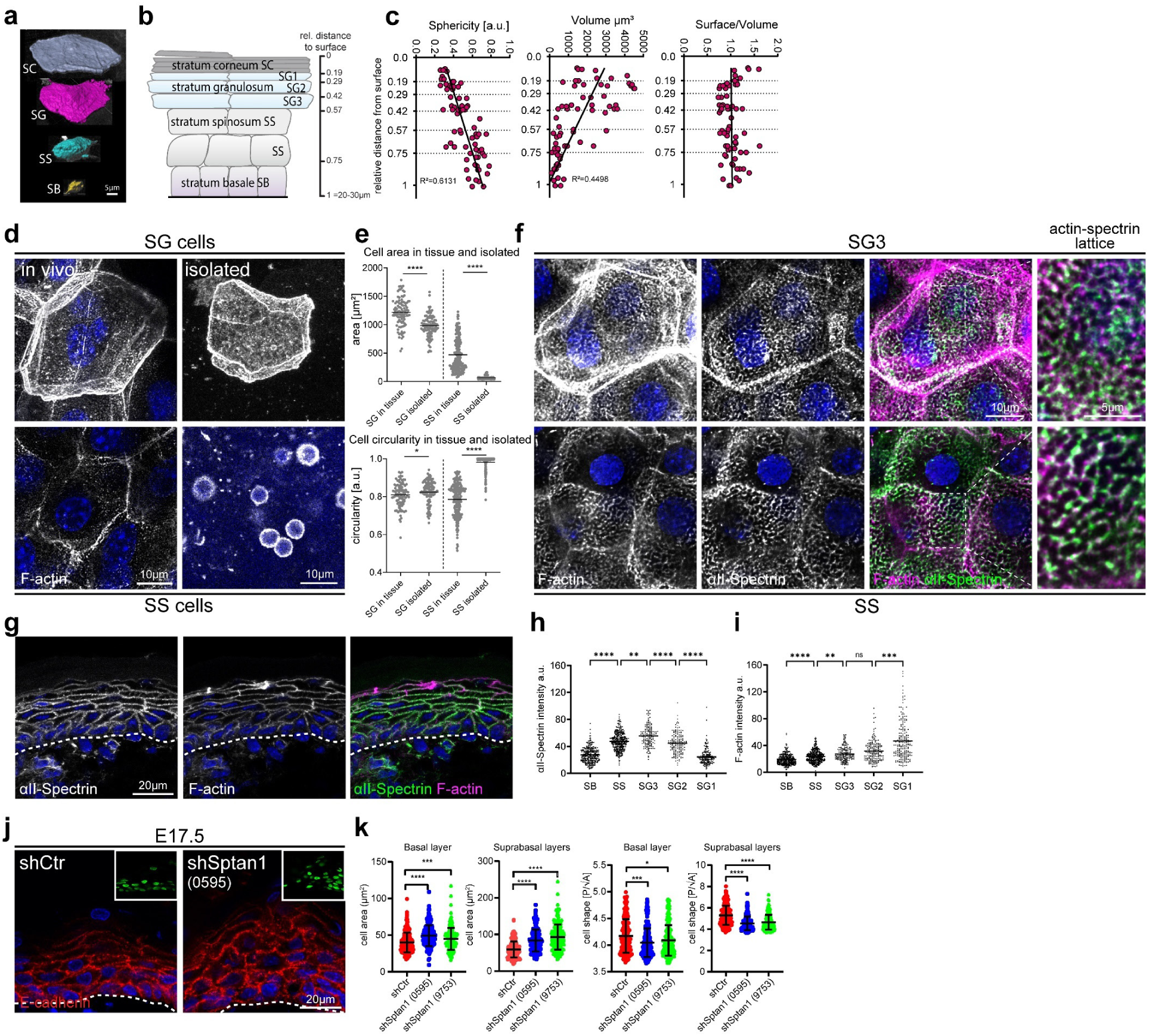
αII-spectrin determines epidermal cell shape. **a** Representative examples of 3D-rendered keratinocytes in the different epidermal layers of newborn epidermal confetti mice using cytoplasmic YFP expression. **b** Schematic illustration and denotation of the epidermal layers and the distance of the boarder of each layer relative to the stratum corneum surface. **c** Quantification of cell shape parameters from rendered cells. 71 rendered cells from 3 mice have been analyzed. R-squared values for the linear regression analysis of the accumulated values are shown. **d** Newborn epidermal whole-mount immunofluorescence analysis for Phalloidin (F-actin) revealing top-view cell shapes in the spinous (SS cells) and granular layer (SG cells) within tissue (in vivo). Right column: Phalloidin staining of single cells isolated from the spinous or granular layer showing deformation upon isolation only in spinous cells. **e** Quantification of cell top view area/shape of granular and spinous cells in the tissue and after isolation using stainings as shown in **d**. \**P* < 0.05, \*\*\*\**P* < 0.0001; >100 cells from 3 mice (isolated) or 6 mice (in tissue) with Kolmogorov-Smirnov per layer. **f** Newborn epidermal whole-mount immunofluorescence analysis for Phalloidin (F-actin) and αII-spectrin. Minimal max. projections of the spinous (SS) and granular layer3 (SG3) are shown. **d,f** representative images of N≥3 biological replicates. **g** Immunofluorescence analysis for Phalloidin (F-actin) and αII-spectrin on newborn skin cryo sections. Representative image of n>3 biological replicates. **h,i** Quantification of cortical αII-Spectrin and F-actin intensity across the epidermal layers. Mean values (lines) of pooled cells (dots) from 3 mice. \*\*\*\**P* < 0.0001, \*\*\**P* = 0.0001, \*\**P* < 0.007; n≥157 cells with Kruskal–Wallis, Dunn’s multiple comparison test. **j** Skin sections from *shCtr* and *shSptan1 0595* transduced E17.5 embryos immunolabeled for E-cadherin. Upper Insets show transduced cells (H2B−GFP+). Dashed lines indicate the dermal-epidermal border. **k** Quantification of cell sagittal area and cell shape (perimeter/√area) from data shown in **j**. Mean (lines) ± SD from ∼200 individual cells (dots) from n=3 embryos per condition. \*\*\*\**P*<0.0001 by Kolmogorov-Smirnov test for sagittal cell area. \*\*\**P* = 0.0002 basal *shSptan1 0595;* *P=0.0181 for basal *shSptan1 9753*, \*\*\*\**P*<0.0001 by Kolmogorov-Smirnov test for cell shape index.

Intercellular AJs are key determinants of cell shape (Luxenburg and Zaidel-Bar, 2019; Niessen et al., 2024). We thus asked whether epidermal cell shape depends on cell-cell contacts. To this end, we adapted a method (Matsui et al., 2021) to separate the suprabasal spinous and first granular layers (SS/SG3) from the late-differentiated (SG2/SG1) layers, to then isolate individual cells. While individual SS/SG3 cells round up upon loss of cell-cell contacts, SG2/SG1 cells maintained their characteristic flattened tetrakaidecahedron shape (Fig. 1d,e; Supplementary Fig. 1a)(Yokouchi et al., 2016a).

The ability of single SG1/SG2 cells to maintain their in vivo shape is reminiscent of red blood cells, in which the spectrin cortical cytoskeleton is essential to preserve their characteristic shape (Leterrier and Pullarkat, 2022). Immunostaining for non-erythrocyte αII-spectrin and F-actin in epidermal whole mounts and sections from newborn mice revealed that spectrin partially colocalized with F-actin to form cortical micro honeycomb-like lattices in suprabasal layers (Fig. 1f,g; Supplementary Fig. 1c). Whereas αII-spectrin was cortically most enriched in the SG3 layer (Fig. 1h), cortical F-actin was increasingly enriched in the upper layers (Rübsam et al., 2017), with the highest enrichment in SG1 cells (Fig. 1i). Thus, during stratification transitions in cell shape are accompanied by changes in cortical actin-to-spectrin ratios, suggesting these changes may induce these shape change. In agreement, suprabasal αII-spectrin enrichment correlated with the initial flattening of suprabasal cells at embryonic (E) day 15 (Supplementary Fig. 1d, arrow), that precedes stratum corneum formation (Rübsam et al., 2017). Thus, spectrin is an integral part of the keratinocyte actin network.

To determine whether αII-spectrin regulates epidermal cell shape changes, we depleted αII-spectrin during epidermal development using lentiviruses encoding two different *Sptan1* short hairpins *(shSptan1-0595 or shSptan1-9753)* or a control *shRNA* (*shScr*) together with a GFP-tagged histone 2B reporter (H2B-GFP) to identify transduced cells (Supplementary Fig. 1e-g)(Beronja et al., 2010). We then quantified the cell area and the cell shape index (perimeter [P]/√area [A]) (Sahu et al., 2020b) as a readout for differentiation-dependent changes in cell shape. Depletion of αII-spectrin resulted in less-flattened suprabasal cells as indicated by an increased cell area and a decrease in shape index (Fig. 1h,i). Interestingly, basal spectrin-depleted cells also increased in size and became even more spherical. Inactivation of αII-spectrin in all epidermal cells using Keratin14-Cre (*Sptan1^epi-/-^*)(Supplementary Fig. 1h,i) confirmed changes in cell shape in newborn epidermis (Supplementary Fig. 1j,k). Thus, αII-spectrin regulates keratinocyte shape in all layers of the epidermis.

**Supplementary Figure 1.**
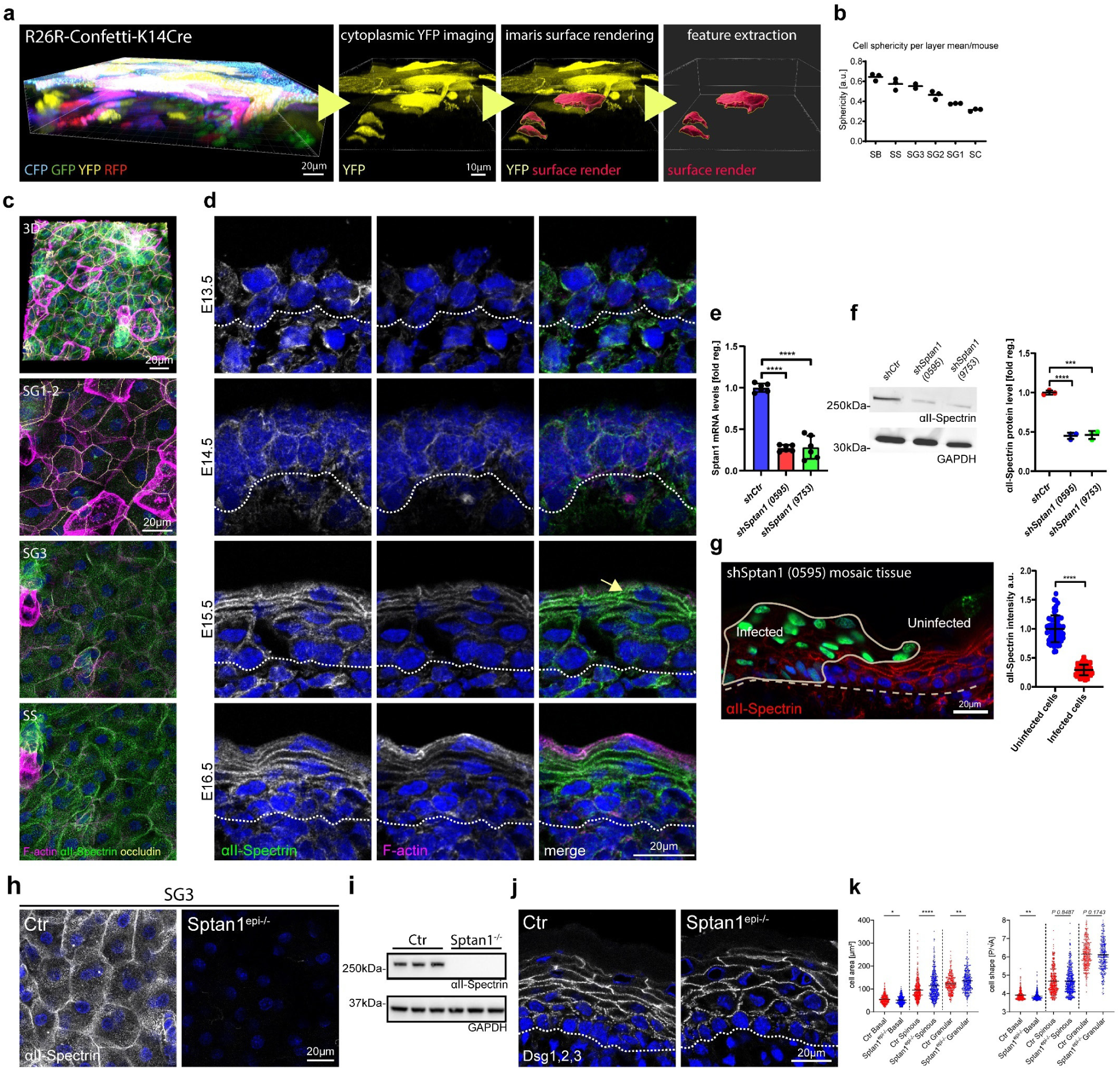
αII-spectrin determines epidermal cell shape. **a** From left to right: 3D whole mount of confetti epidermis from newborn mice showing expression of all 4 transgenic confetti colors; imaging of the cytoplasmic YFP signal; overlay of the rendering based on the YFP signal; rendered cell volumes. **b** Quantification of cell sphericity from rendered cells per layer corresponding to Fig. 1c. Dots: mean values per mouse. **c** Newborn epidermal whole-mount immunofluorescence analysis for Phalloidin (F-actin), αII-spectrin and tight junction marker occludin marking the SG2 layer. Overview of protein distribution across layer corresponding to Fig. 1c. Minimal max. projections of the epidermal layers and a full projection (3D) are shown. **d** Immunofluorescence analysis for Phalloidin (F-actin) and αII-spectrin on newborn epidermis cryo sections of embryonic timepoints as indicated. Dashed line marks the epidermal-dermal boarder. **c,d** representative images of N≥3 biological replicates. **e** Quantitative qPCR analysis of Sptan1 mRNA in primary mouse keratinocytes transduced with Ctr shRNA or one of two *Sptan1* -specific shRNAs (0595 and 9753). Mean ± SD of six preparations. \*\*\*\**P* <0.0001 by unpaired t-test. **f** Western blot analysis of primary mouse keratinocytes transduced with *Scr*, *Sptan1 0595*, or *Sptan1 9753* shRNAs and quantification of αII-spectrin protein levels. Data are the mean ± SD of three preparations. **** *P*>0.0001, \*\*\**P*=0.0006 by unpaired t-test. **g** Dorsal skin mosaic tissue sections from *shSptan1 0595* transduced E17.5 embryos immunolabeled for αII-spectrin. Line indicates areas of infected cells; dashed line indicates the dermal-epidermal border. Nuclei were stained with DAPI. Quantification of αII-spectrin intensity. Data are the mean ± SD of 60 individual cells from n=3 embryos. Bars: mean normalized intensity; dots: individual cells. \*\*\*\**P*>0.0001 by unpaired t-test. **h** Newborn epidermal whole-mount immunofluorescence analysis αII-spectrin in Ctr and αII-spectrin deficient epidermis (*Sptan1^epi-/-^*). Max. projection of the SG3 layer. **i** Western blot analysis of primary mouse keratinocytes isolated from Ctr and αII-spectrin deficient epidermis (*Sptan1^epi-/-^*). **j** Immunofluorescence analysis for shape using combined staining for desmoglein1,2,3 (Dsg1,2,3) on Ctr and αII-Spectrin deficient newborn epidermis sections. **k** Quantification of cell sagittal area and shape (perimeter/√area)/layer using stainings as shown in **j**. \*\*\*\**P* < 0.0001, \*\**P* < 0.005, \**P* < 0.05; cells: n=701 (basal), n=927 (spinous), 688 (granular) with Kolmogorov-Smirnov per layer.

### E-cadherin controls cell shape upstream of spectrin

We next asked how spectrin is recruited to the cortex in keratinocytes. One candidate is ankyrin, a major binding partner of spectrin that enables spectrin self-assembly into networks (Bennett and Healy, 2009; Bennett and Lorenzo, 2016), and also interacts with E-cadherin (Kong et al., 2023; Kizhatil et al., 2007; Bennett and Lorenzo, 2013). Immunostaining of E17.5 embryos for ankyrin G (encoded by *Ank3*), which is expressed in the developing epidermis (Sennett et al., 2015), revealed cortical enrichment in the granular layer that is lost upon spectrin depletion (Fig. 2a,c,d). In contrast, *Ank3* depletion in E17.5 embryos (Supplementary Fig. 2a-c) did not alter spectrin localization nor levels (Fig. 2a, b), with also no changes in F-actin nor cell shape (Supplementary Fig. 2d-g).

**Figure 2.**
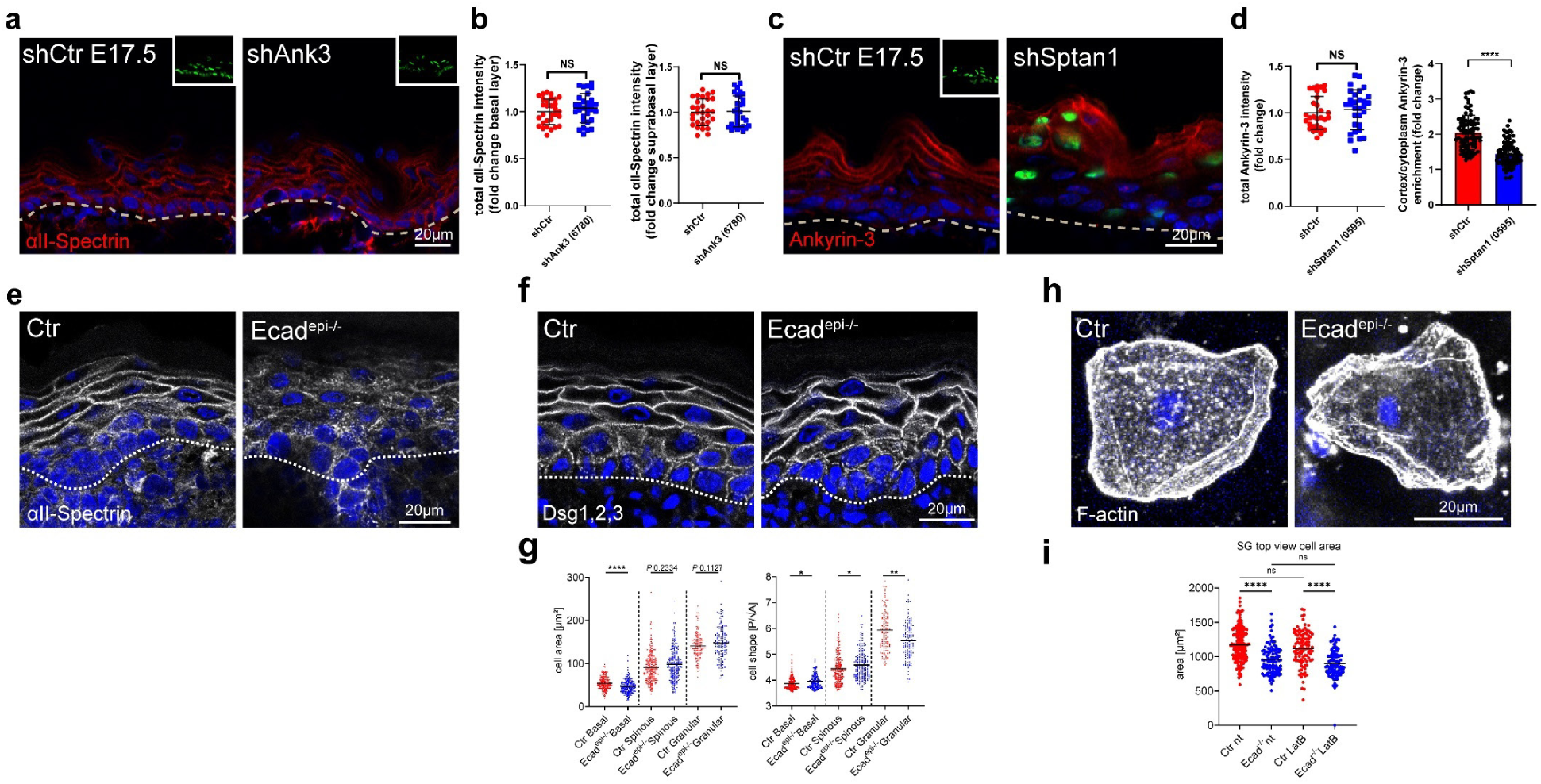
E-cadherin controls cell shape upstream of spectrin. **a** Sagittal views of dorsal skin sections from *shCtr* and shAnk3 6780 transduced E17.5 embryos immunolabeled for αII-spectrin. Insets show the transduced cells (H2B−GFP+). **b** Quantification of basal (left) and suprabasal (right) layer αII-spectrin intensity from data shown in **a**. Mean ± SD of 30 ROI from n=3 embryos per condition. Bars: mean normalized intensity; dots: microscopy fields. NS: *P*=0.3199 (basal), NS: *P*=0.8256 by unpaired t-test. **c** Dorsal skin sections from *shCtr* and mosaic *shSptan1* 0595 transduced E17.5 embryos immunolabeled for Ankyrin-3. Upper insets show the transduced cells (H2B−GFP+). **d** Left graph: Quantification of Ankyrin-3 intensity from data shown in **c**. Mean ± SD of 30 ROI from n=3 embryos per condition. Bars: mean normalized intensity; dots: microscopy fields. NS: *P*=0.512 by unpaired t-test. Right graph: Quantification of Ankyrin-3 cortical enrichment from the data shown in **c**. Mean ± SD from 90 individual cells from n=3 embryos per condition. Bars: Ankyrin-3 cortex/cytoplasm intensity ration mean; dots: individual cells. \*\*\*\**P* <0.0001 by unpaired t-test. **e** αII-Spectrin staining on Ctr and E-cadherin deficient newborn epidermis sections showing impaired cortical recruitment of αII-Spectrin upon loss of E-cadherin. Representative images of n=3 biological replicates. **f** Immunofluorescence analysis for shape using combined staining for desmoglein1,2,3 (Dsg1,2,3) on Ctr and E-cadherin deficient newborn epidermis sections. **g** Quantification of cell sagittal area and shape (perimeter/√area)/layer using stainings as shown in **f**. \*\**P* < 0.005, \**P* < 0.03; cells: n=420 (basal), n=445 (spinous), 277 (granular) with Kolmogorov-Smirnov per layer. **h** Phalloidin staining of single cells isolated from the granular layer of Ctr or E-cadherin deficient newborn epidermis. **i** Quantification of the cell topview area of isolated SG cells from Ctr and E-cadherin deficient newborn epidermis as shown in **h** and treated with Latrunculin B (0.1µM). Dots represent individual cells isolated from three mice. \*\*\*\**P* < 0.0001; n≥98 cells with Kruskal–Wallis, Dunn’s multiple comparison test. All images: Nuclei were stained with DAPI (blue).

E-cadherin-based AJ can recruit spectrin to the cortex (Pradhan et al., 2001). Epidermal inactivation of E-cadherin (*Ecad^epi-/-^)* (Tunggal et al., 2005) resulted in a loss of αll-spectrin from the cortex of suprabasal layers (Fig. 2e), whereas F-actin remained cortical with increased intensity in the spinous layer (Rübsam et al., 2017). In contrast, E-cadherin maintained its localization in αII-spectrin-depleted epidermis (Fig. 1h). Consistently, αII-spectrin was mislocalized upon depletion of E-cadherin in cultured keratinocytes (Supplementary Fig. 1h), whereas depletion of αII-spectrin did not affect AJ assembly (Supplementary Fig. 1i).

Using *Ecad^epi-/-^* mice we then asked whether E-cadherin-dependent recruitment of spectrin controls cell shape. Epidermal loss of E-cadherin altered cell shapes in all layers with a reduced flattening of SG cells, as seen with spectrin depletion (Fig. 2f, g). Interestingly and in contrast to spectrin depletion, spinous layer cells showed increased flattening upon loss of E-cadherin, perhaps as a result of increased cortical actin in this layer. Isolated E-cad^-/-^ SG1/SG2 cells were also smaller but maintained their flattened tetrakaidecahedron cell shape (Fig. 2h, i). Moreover, latrunculin B treatment of isolated SG2/SG3 did not alter their flattened tetrakaidecahedron shape (Fig. 2h, i), despite efficient actin depolymerization (Supplementary Fig. 2j, k). Together, these data indicate that E-cadherin through polarized organization of the spectrin-actomyosin cortex regulates transitions in cell shape that guide terminal differentiation when cells move up through the epidermis with the shape of SG2 cells once set becoming independent of AJ or actomyosin.

**Supplementary Figure 2.**
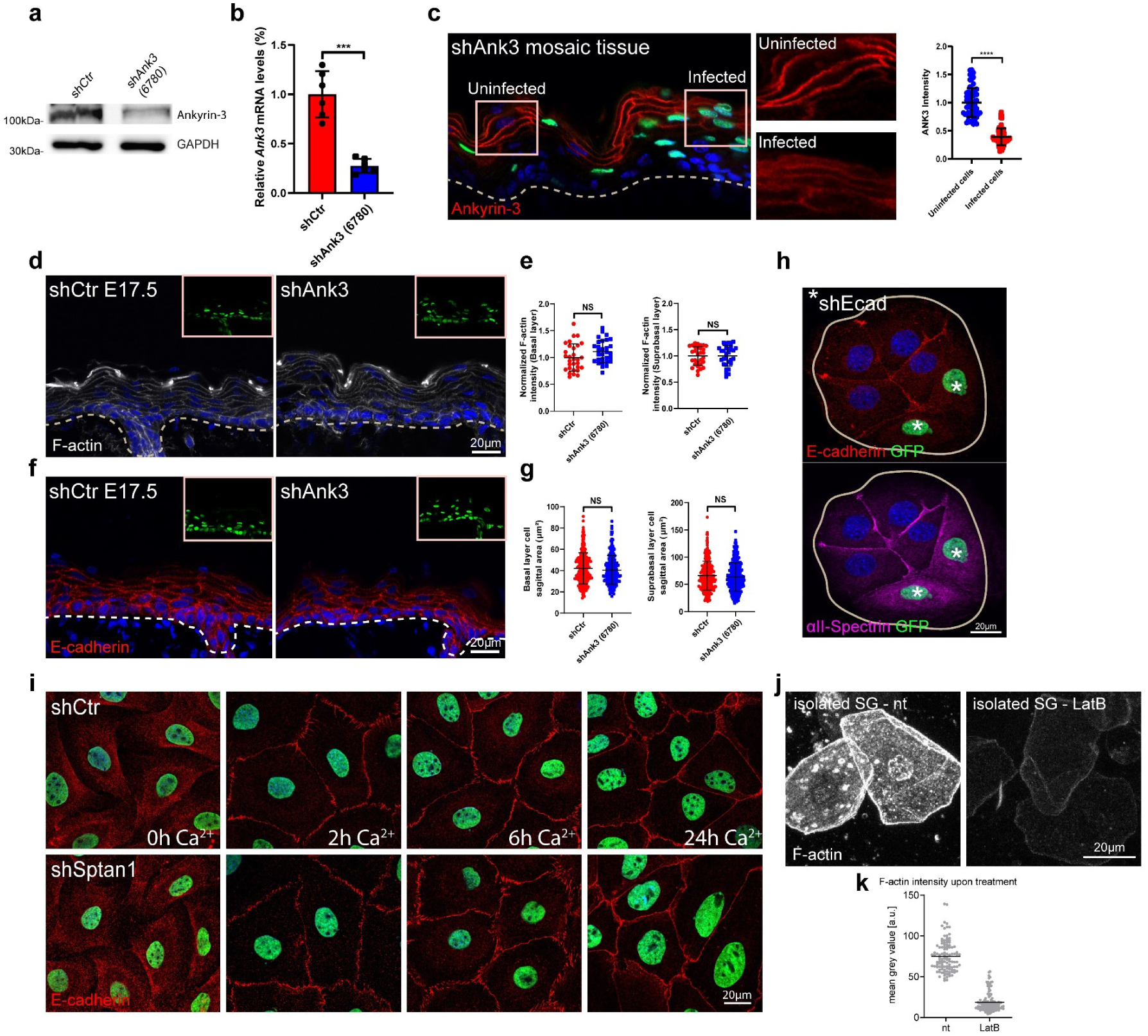
E-cadherin controls cell shape upstream of spectrin. **a** Western blot analysis of primary mouse keratinocytes transduced with *shCtr* and Ank3 6780 shRNAs. Blots were probed for Ankyrin-3 and GAPDH. **b** qPCR analysis of Ank3 mRNA in primary mouse keratinocytes transduced with *shCtr* shRNA or Ank3 specific shRNAs (6780). Data are the mean ± SD of six preparations. \*\*\**P*=0.0004 by unpaired t-test. **c** sagittal views of dorsal skin mosaic tissue sections from shAnk3 6780 transduced E17.5 embryos immunolabeled for Ankyrin-3. Graph: Quantification of Ankyrin-3 intensity. Mean ± SD of 60 individual cells from n=3 embryos. Bars: mean normalized intensity; dots: individual cells. \*\*\*\**P*>0.0001 by unpaired t-test. **d** Sagittal views of dorsal skin sections from *shCtr* and shAnk3 6780 transduced E17.5 embryos immunolabeled for F-actin. Insets show transduced cells (H2B−GFP+). **e** Quantification of basal (left) and suprabasal (right) layer F-actin intensity from data shown in **d**. Data are the mean ± SD of 30 ROI (Region of interest) from n=3 embryos per condition. Horizontal bars represent the mean normalized intensity, and circles/squares represent microscopy fields. NS: *P*=0.0692 (basal), NS: *P*=0.9506 (suprabasal) by unpaired t-test. **f** Sagittal views of dorsal skin sections from *shCtr* and shAnk3 6780 transduced E17.5 embryos immunolabeled for E-cadherin. Insets show transduced cells (H2B−GFP+). **g** Quantification of basal (left) and suprabasal (right) layer cell cross-section area from data shown in **f**. Data are the mean ± SD from∼ 200 individual cells from n=3 embryos per condition. Horizontal bars represent the cell cross-section area mean, and circles/squares represent individual cells. NS: *P*=0.2220 (basal), NS: *P*=0.796 (suprabasal) by Kolmogorov-Smirnov. **h** Immunofluorescence analysis for the recruitment of E-cadherin and αII-spectrin to intercellular contacts in *shCtr* and *shEcad* 2287 transduced primary keratinocytes (GFP positive nuclei, asterisks). Representative images of n=3 biological replicates each. **i** Immunofluorescence analysis for E-cadherin (red) based early intercellular contacts in *shCtr* and *shSptan1 0595* transduced primary keratinocytes at indicated Ca^2+^ timepoints. Representative images of n=3 biological replicates each. **j** Phalloidin staining of single cells isolated from granular layer treated with or without (nt) Latrunculin B (0.1µM). **k** Quantification of F-actin intensity (Phalloidin) of isolated SG cells as shown in **j** treated with or without Latrunculin B. Dots represent individual cells pooled from three mice.

### Cortical F-actin and spectrin organization are mutually dependent

We next asked whether F-actin and spectrin reciprocally coordinate the organization of the cortex into a honeycomb lattice. Treatment of E17.5 embryos with low levels of Latrunculin B to reduce F-actin but not disturbing E-cadherin adhesion (Supplementary Fig. 3a) reduced cortical αII-spectrin levels in all layers of the epidermis (Fig. 3a, b). Conversely, embryonic KD of αII-spectrin reduced suprabasal cortical F-actin levels of E17.5 embryos (Fig. 3c, d). High-resolution imaging of newborn epidermal whole mounts revealed a striking loss of F-actin honeycomb lattices in *Sptan1^epi-/-^* mice, with over 60% of suprabasal cells showing a more streak- or spot-like organization (Fig. 3e,f, arrow), in contrast to 90% of control cells showing honeycomb lattices, indicating that spectrin not only recruits but also organizes F-actin at the cortex.

**Figure 3.**
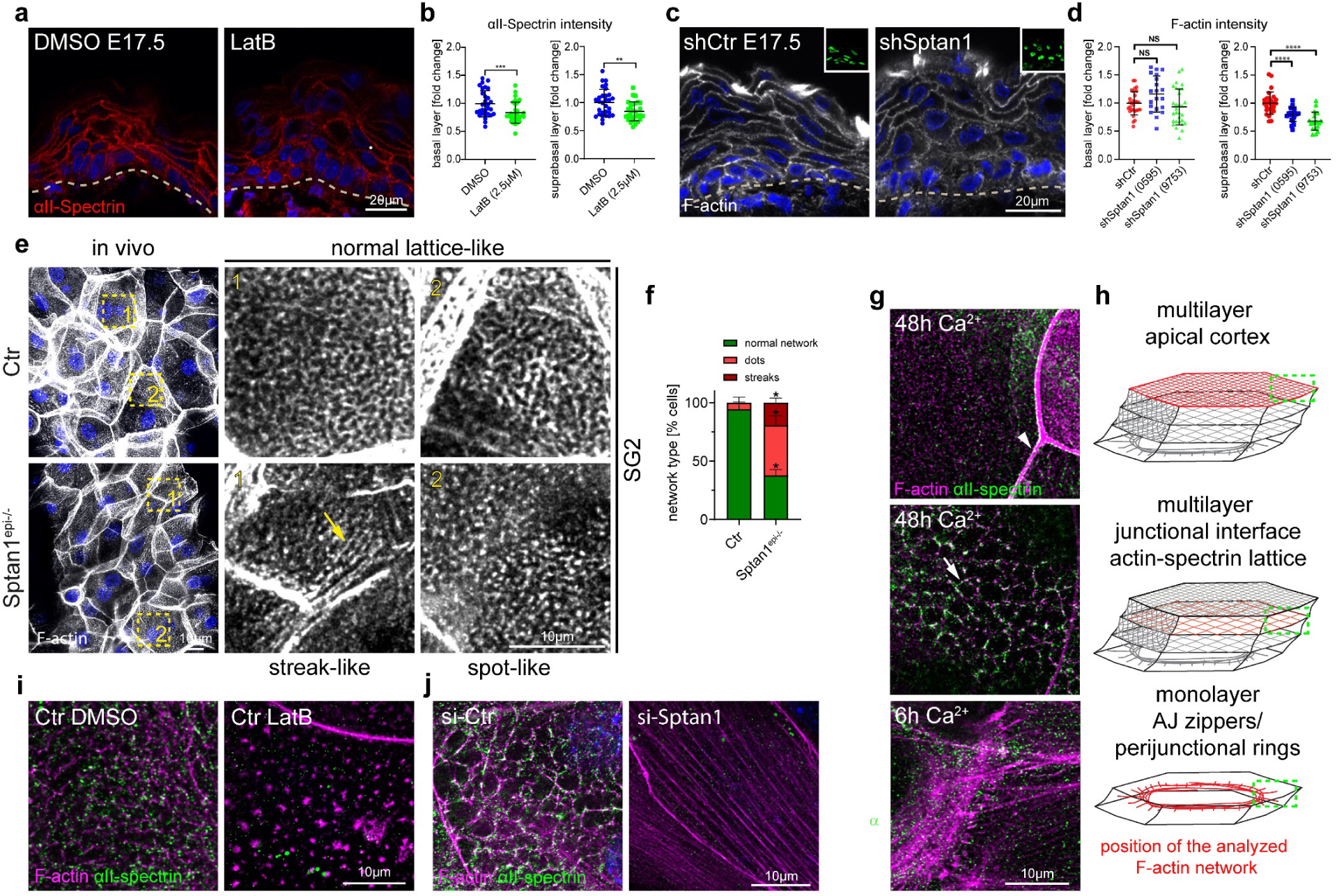
Cortical F-actin and spectrin organization are mutually dependent. **a** Dorsal skin sections from E17.5 wild-type embryos treated with DMSO or latrunculin B (2.5µM) immunolabeled for αII-spectrin. **b** Quantification of basal (left graph) and suprabasal (right graph) layer αII-spectrin intensity from data shown in **a**. Mean ± SD of 30 ROI from n=3 embryos per condition. Bars: mean normalized intensity; dots: microscopy fields. \*\*\**P*=0.005 (basal, latrunculin B), \*\**P*=0.0044 (suprabasal, latrunculin B) by unpaired t-test. **c** Dorsal skin sections from *shCtr* and mosaic *shSptan1* 0595 transduced E17.5 embryos immunolabeled for F-actin. Upper insets show the transduced cells (H2B−GFP+). **d** Quantification of basal (left) and suprabasal (right) layer F-actin intensity from data shown in **c**. Mean ± SD of 30 ROI from n=3 embryos per condition. Bars: mean normalized intensity; dots: individual microscopy fields. NS: P=0.0553 (*shSptan1* 0595, basal), P=0.2818 (*shSptan1* 9753, basal), ****P=0.0004 (*shSptan1* 0595, suprabasal), ****P=0.0006 (*shSptan1* 9753, suprabasal) by unpaired t-test. Nuclei were stained with DAPI, dotted lines indicate the dermal-epidermal border. NS, not significant. **e** Newborn epidermal whole-mount immunofluorescence analysis for cortical F-actin organization upon loss of αII-spectrin. Note streak-like (arrow) and spot-like reorganization of F-actin upon loss of αII-spectrin. Representative images of 4 biological replicates (mice). **f** Quantification of the percentage of cells in the granular layer showing either normal (lattice-like) or abnormal (streak-like or spot-like) F-actin organization. *P<0.05 with Mann-Whitney for the mean of n=4 biological replicates including 466 (Ctr) and 385 (Sptan1KO) analyzed cells. **g** Immunofluorescence analysis for αII-Spectrin and F-actin after 6h or 48h in high Ca^2+^ at cell-cell interfaces (arrow: actin-spectrin lattice, arrowhead: TJ-supporting apical F-actin ring) and apical surface. Representative images of n≥3 biological replicates each. **h** Schematic representation of the image position in the mono or multilayered keratinocytes. **i** Immunofluorescence analysis for αII-spectrin and F-actin after 48h in high Ca^2+^ at cell-cell interfaces with and without Latrunculin B treatment (1h, 0.1µM). **j** Immunofluorescence analysis for αII-spectrin and F-actin (48h high Ca^2+^) upon αII-spectrin knockdown. Representative images of n=3 biological replicates each.

We then asked how AJ recruit and organize the actin-spectrin network by switching primary keratinocytes to high Ca^2+^ (1.8 mM) to induce contact formation and stratification. Upon initial E-cadherin engagement (6h Ca^2+^) only F-actin but not spectrin was recruited to early AJ, so-called AJ zippers (Fig. 3g, h), to form peri-junctional rings (Vasioukhin et al., 2000) (Fig. 3g, h; Supplementary Fig. 3b). Consistently, αII-spectrin knockdown did not affect initial E-cadherin engagement or actin recruitment (6h Ca^2+^, Supplementary Fig. 2i). Stratification of keratinocytes into a multilayer (48h Ca^2+^) upregulated spectrin protein levels (Supplementary Fig. 3c, 2.28-fold compared to 6h Ca^2+^), in agreement with the *in vivo* suprabasal increase in cortical spectrin (Fig. 1g, h). Importantly, as seen in vivo, spectrin partially co-localized with F-actin and E-cadherin to form micro-honeycomb-like cortical F-actin-spectrin lattices, (Fig. 3g, arrow, h; Supplementary Fig. 3b, d). Spectrin was also localized to the F-actin cortical ring that supported barrier-forming TJs in the uppermost apical cell layer, albeit to a much lesser extent than F-actin (Fig. 3g, h, arrowhead; Supplementary Fig. 3b)(Rübsam et al., 2017). In contrast, little αII-spectrin was recruited to the F-actin cortex supporting the apical membrane that does not form cell-cell contacts (Fig. 3g, h; Supplementary Fig. 3b), in agreement that E-cadherin is essential to recruit αII-spectrin to the cortex. Low levels of Latrunculin B reduced cortical αII-spectrin recruitment (Fig. 3i, Supplementary Fig. 3e), similar to in vivo. Strikingly, depletion of αII-spectrin resulted in a loss of cortical honeycomb lattices, with F-actin showing a linear fiber organization reminiscent of F-actin stress fibers (Fig. 3j). In contrast, αII-spectrin KD did not prevent formation of the apical TJ-associated F-actin cortical ring even though F-actin levels were reduced (Supplementary Fig. 3h, i).

Thus, F-actin and spectrin cortical localization and organization are interdependent. Upon initial adhesion, AJs first recruit F-actin to then reorganize the F-actin cortex (Vaezi et al., 2002) allowing recruitment of αII-spectrin that then is necessary to organize F-actin-spectrin lattices in suprabasal layers.

**Supplementary Figure 3.**
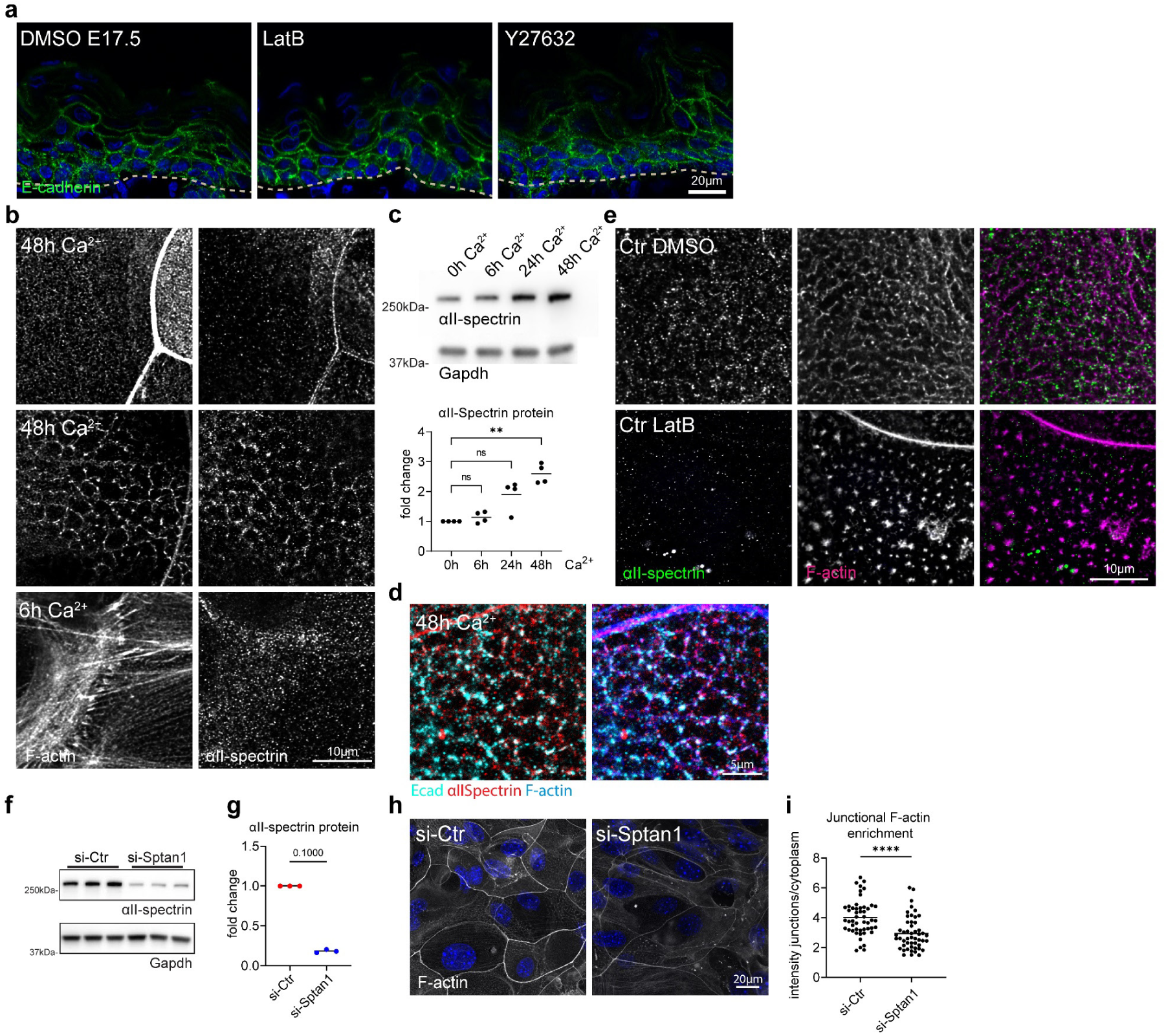
Cortical F-actin and spectrin organization are mutually dependent. **a** Sagittal views of dorsal skin sections from E17.5 wild-type embryos treated with DMSO, latrunculin, and Y27632 immunolabeled for E-cadherin. Nuclei were stained with DAPI; dotted lines indicate the dermal-epidermal border. NS, not significant. **b** Immunofluorescence analysis for αII-Spectrin and F-actin after 6h or 48h in high Ca^2+^ at cell-cell interfaces and apical surface. Single channels corresponding to Fig. 3g. **c** Western blot analysis and quantification for αII-spectrin protein levels after Ca^2+^ switch for the timepoint indicated. Normalized to GAPDH and to 0h Ca^2+^ timepoint. Dots represent biological replicates from n=4 independent primary keratinocyte isolates. \*\**P*=0.003 with Kruskal–Wallis, Dunn’s multiple comparison test. **d** Immunofluorescence analysis for E-cadherin, αII-spectrin and F-actin after 48h in high Ca^2+^ at cell-cell interfaces **e** Immunofluorescence analysis for αII-spectrin and F-actin after 48h in high Ca^2+^ at cell-cell interfaces with and without Latrunculin B treatment (1h, 0.1µM). Single channels corresponding to Fig. 3i. **f** Western blot analysis for αII-spectrin protein levels upon siRNA (siPools) mediated knockdown 96h post transfection (72h Ca^2+^). **g** Western blot quantification for αII-spectrin as shown in **f**, normalized to GAPDH. Dots represent biological replicates, n=3 with Mann-Whitney. Representative example of n=6 independent primary keratinocyte isolates. **h** Immunofluorescence analysis of F-actin organization at apical junction rings after 48h in high Ca^2+^ and siRNA mediated knockdown of αII-spectrin. **i** Quantification of F-actin intensity at apical junction rings (mean grey value, junctions/cytoplasm) as shown in **h**. Dots represent pooled values of single cells from n=3 biological replicates. \*\*\*\**P*<0.0001 with Kolmogorov-Smirnov. Right graph: Mean values from n=3 biological replicates tested with Mann-Whitney.

### Spectrin stabilizes contractile cortical actomyosin networks

The cortex of epidermal keratinocytes is highly enriched for the actin-binding protein myosin II that regulates contractile states of the actin cytoskeleton (Dor-On et al., 2017; Sumigray et al., 2012). In *Drosophila*, loss of either β- or α-spectrin increased junctional myosin levels and activity (Ibar et al., 2023; Deng et al., 2015; Forest et al., 2018). The observed reorganization of F-actin into stress fiber-like structures upon loss of αII-spectrin thus suggested a change in myosin recruitment and activity. Staining for non-muscle myosin heavy chain IIa (myosin-IIa) revealed that myosin-IIa was enriched at lateral junctions in the spinous layer whereas in the granular layer it was enriched in spots that decorated the actin-spectrin honeycomb lattices, thus indicating an increase in its tensile state as cells moved from the spinous to the granular layer. Loss of spectrin increased myosin intensity and spot size, suggesting an increase in the contractile state of F-actin networks (Fig. 4a), similar to Drosophila. Vice versa, inhibition of myosin contractility using the Rho-associated protein kinase (ROCK) inhibitor, Y27632, decreased cortical αII-spectrin levels in E17.5 embryos (Fig. 4b, c), thus indicating that spectrin and myosin II jointly coordinate organization and tensile states of the F-actin cortex

**Figure 4.**
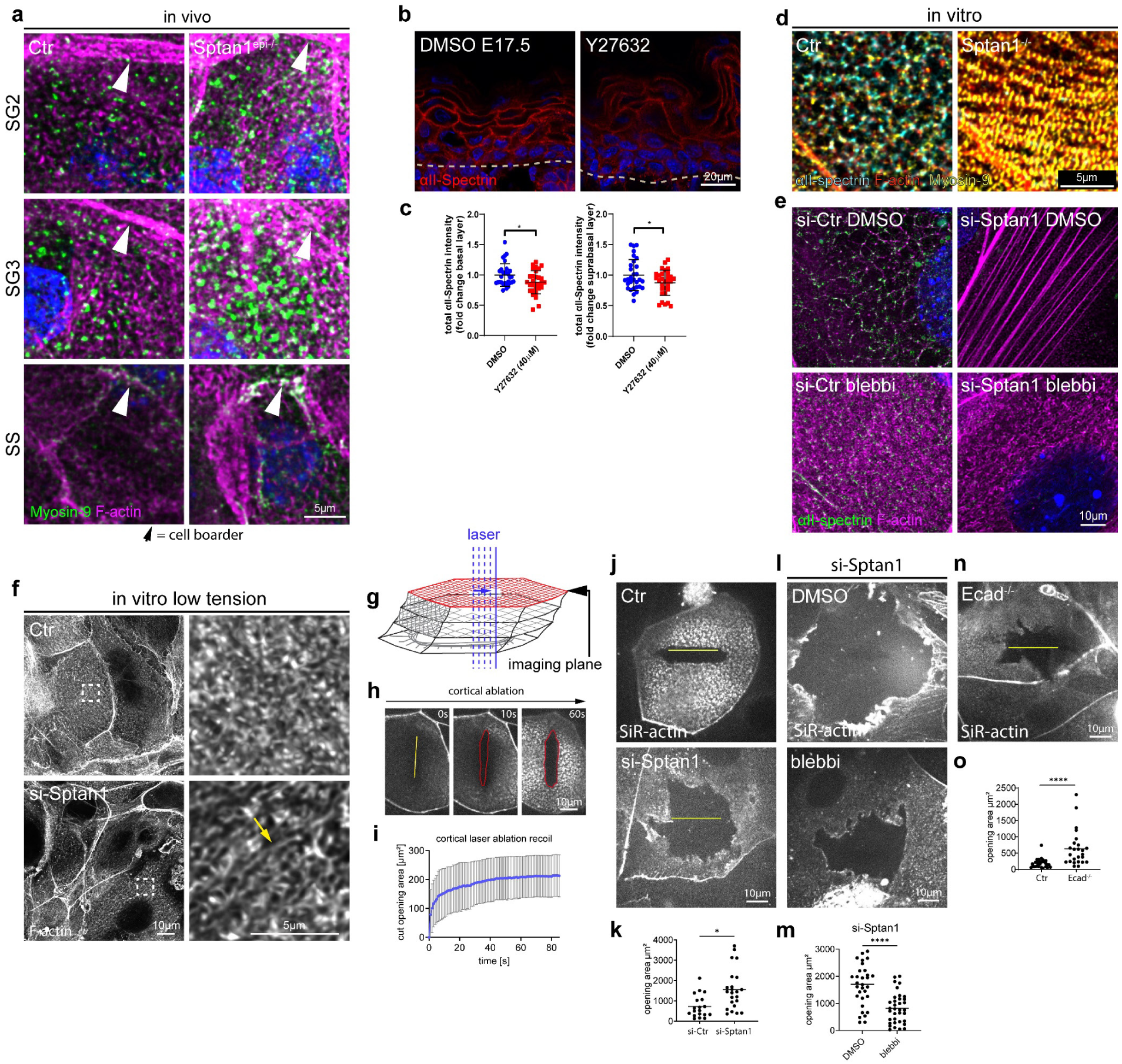
Spectrin determines junctional actomyosin network structure and stability. **a** Newborn epidermal whole-mount immunofluorescence analysis for Phalloidin (F-actin) and non-muscle myosin heavy chain IIa (Myosin-IIa). Minimal max. projections of the denoted layers are shown. **b** Dorsal skin sections from E17.5 wild-type embryos treated with DMSO or Y27632 (40µM) immunolabeled for αII-spectrin. **c** Quantification of basal (left graph) and suprabasal (right graph) layer αII-spectrin intensity from data shown in **b**. Data are the mean ± SD of 30 ROI from n=3 embryos per condition. Bars: mean normalized intensity; dots: microscopy fields. \**P*=<0.05 by unpaired t-test. **d** Immunofluorescence analysis for αII-spectrin, F-actin and non-muscle myosin heavy chain IIa (Myosin-IIa) (48h high Ca^2+^) in Ctr and αII-spectrin deficient (KO) cells. **e** Immunofluorescence analysis for αII-spectrin and F-actin (48h high Ca^2+^) upon αII-spectrin (*Sptan1*) knockdown and treatment with either DMSO or low dose blebbistatin (5µM). Representative images of n=3 biological replicates each. **f** Immunofluorescence analysis for F-actin (48h high Ca^2+^) upon αII-spectrin (*Sptan1*) knockdown at low tension (low dose blebbistatin 5µM) showing streak like defects similar to in vivo. Representative images of n=3 biological replicates each. **g** Illustration of laser ablation in multilayered keratinocytes. **h** Laser ablation: Still images from live imaging of SiR-actin (F-actin) labeled Ctr cells (48h high Ca^2+^) at indicated timepoints after 17µm line ablation showing progressive elliptical cortical openings. **i** quantification of the elliptical opening area (red line) i.e. recoil over time. Line: mean +/-SD opening area of 13 ablations from N=4 biological replicates. **j** Cortical laser ablation: Live imaging of SiR-actin (F-actin) labeled Ctr and αII-spectrin knockdown keratinocytes after 48h high Ca^2+^. Cells are depicted 10 seconds after a linear laser cut (yellow line). The opening in the F-actin cortex is seen as black area around the yellow line. **k** Quantification of opening areas in the apical F-actin cortex upon linear laser ablation as shown in **j**. Lines represent means, dots represent single openings/cells pooled from n=3 independent experiments/biological replicates. \**P*=0.0215 with Kolmogorov-Smirnov. **l** Laser ablation (as described for **j**) of αII-spectrin knockdown keratinocytes treated with DMSO or low dose blebbistatin (5µM) 1h prior to ablation. **m** Quantification of opening areas in the apical F-actin cortex upon linear laser ablation as shown in **l**. Lines represent means, dots represent single openings/cells pooled from n=3 independent experiments/biological replicates. \*\*\*\**P*<0.0001 with Kolmogorov-Smirnov. **n** Laser ablation (as described for **j**) of E-cadherin^-/-^ keratinocytes after 48h high Ca^2+^. **o** Quantification of opening areas in the apical F-actin cortex upon linear laser ablation as shown in **n**. Lines represent means, dots represent single openings/cells pooled from n=3 independent experiments/biological replicates. ****P<0.0001 with Kolmogorov-Smirnov.

We next used primary keratinocytes to assess whether and how spectrin-dependent recruitment of myosin II altered the tensile state of the cortex. Co-staining of F-actin, αII-spectrin and Myosin-IIa in stratified control keratinocytes also revealed a spot-like integration of myosin IIa into the spectrin-F-actin lattices, whereas in the absence of αII-spectrin myosin IIa showed an increased and more periodic decoration of cortical F-actin fibers, indicating that these fibers are highly contractile (Fig. 4d; Supplementary Fig. 4a). Lowering myosin II motor activity with blebbistatin (5µM) in control keratinocytes was sufficient to reorganize honeycomb spectrin-F-actin lattices into ultrafine almost diffuse F-actin structures that showed reduced spectrin association (Fig. 4e; Supplementary Fig. 4b), showing that tension is necessary for honeycomb organization. Moreover, blebbistatin treatment disassembled the linear F-actin fibers induced by αII-spectrin depletion, demonstrating that these fibers are indeed under tension. Instead, F-actin was organized into ultrafine F-actin structures similar to those observed in blebbistatin-treated controls except that their appearance was more anisotropic and streak-like (Fig. 4f, arrow; Supplementary Fig. 4b), thus now more resembling the in vivo F-actin defects observed in the αII-spectrin– deficient epidermis (Fig. 3e). Together, these data indicate that loss of αII-spectrin increases tension, but this increase in vivo is lower than that in cultured primary keratinocytes.

We then assessed the mechanical properties of the cortex in the presence of suprabasal spectrin-actomyosin-spectrin lattices by performing linear laser ablation experiments to measure strain-dependent recoil in stratified keratinocytes. Prior to formation of cortical actomyosin-spectrin lattices, the ablation of F-actin-positive intercellular junctions elicited a viscoelastic recoil response of junctional vertices as described for simple epithelial cells (Supplementary Fig. 4c, d) (Wu et al., 2014). In contrast, virtually no recoil of junctional vertices was observed once actomyosin-spectrin lattices were formed (Supplementary Fig. 4c, d). Either subsequent ablation of the lattices itself adjacent to the already-ablated junctions or ablation of only these lattices, induced a strong recoil response with elliptical openings that increased in size over time (Fig. 4g-i; Supplementary Fig. 4e, f). These data indicate that the cortical actomyosin-spectrin lattices exhibit viscoelastic properties as shown previously in zebrafish and C. elegans (Saha et al., 2016; Thi Kim Vuong-Brender et al., 2017). Interestingly, curved intercellular borders straightened upon ablation, indicating that the actomyosin-spectrin lattice regulated the tensional state of AJs to control cell shape (Supplementary Fig. 4g). Importantly, depleting αII-spectrin resulted in a faster, two-fold larger recoil with highly irregular openings (Fig. 4j, k), which was reversed upon blebbistatin treatment. Further, E-cadherin loss induced similar recoil behavior with highly irregular, larger openings (Fig. 4n, o). Thus, E-cadherin-dependent organization or spectrin-dependent lattice organization is necessary to dissipate the myosin-driven increase in tension across the network to prevent damage (Fig. 4l, m). Taken together, these data suggest a model in which E-cadherin through spectrin recruitment dissipates tension to increase the stability of cortical actomyosin lattices under tension.

**Supplementary Figure 4.**
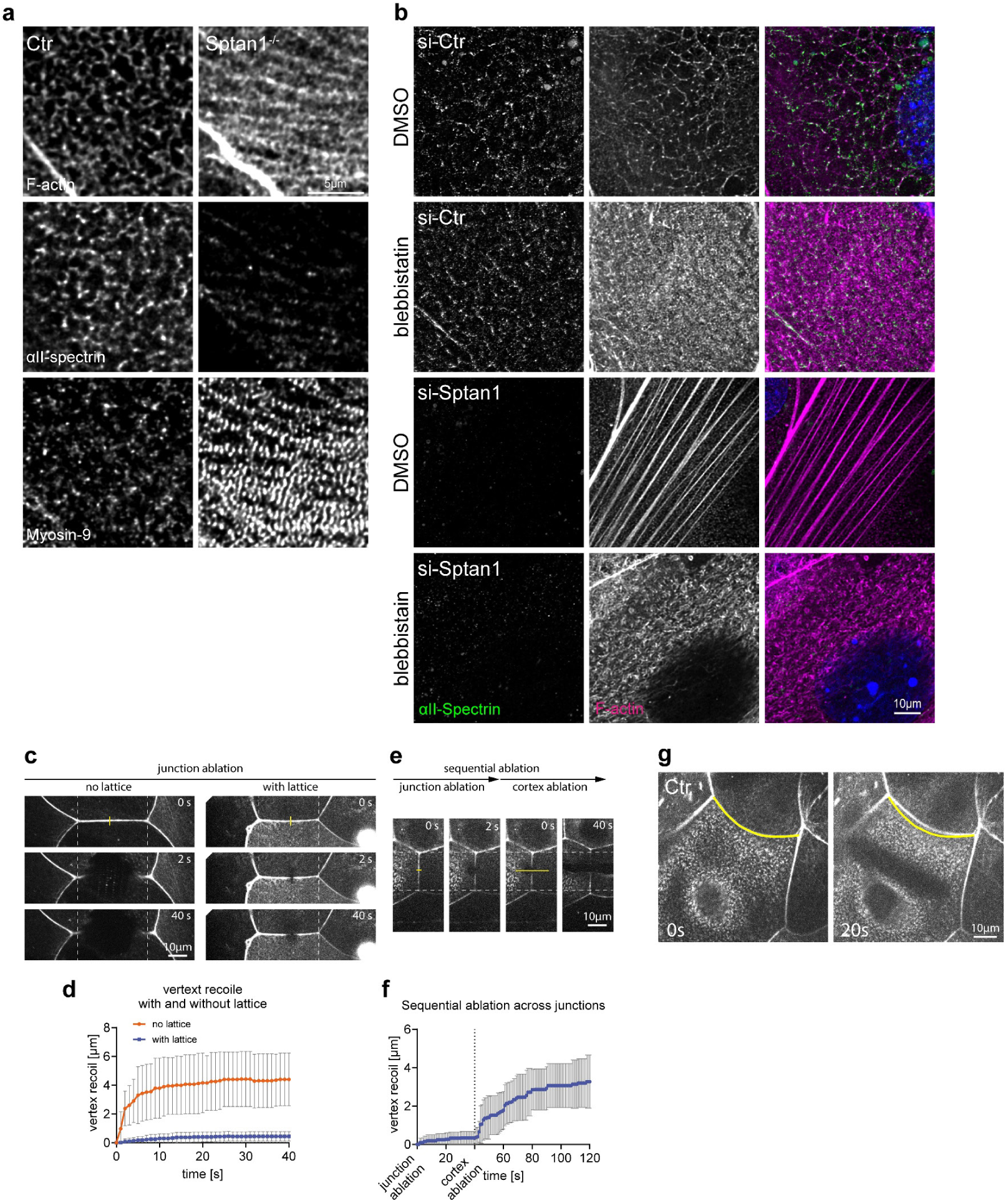
Spectrin determines junctional actomyosin network structure and stability. **A** Immunofluorescence analysis for αII-spectrin, F-actin and non-muscle myosin heavy chain IIa (Myosin-IIa) (48h high Ca^2+^) in Ctr and αII-spectrin deficient (KO) cells. Single channels corresponding to Fig. 4d. **b** Immunofluorescence analysis for αII-spectrin and F-actin (48h high Ca^2+^) upon αII-spectrin knockdown and treatment with either DMSO or low dose blebbistatin (5µM). Representative images of n=3 biological replicates each. Single channel grey scale and merged channels corresponding to Fig. 4e. **c** Junctional laser ablation in apical keratinocytes with (48h Ca^2+^) or without (40h Ca^2+^) a cortical F-actin lattice. Timepoints after ablation (yellow line) are shown as indicated. **d** Quantification of the increase in vertex (dashed line) distance upon ablation. Lines: mean +/-SD recoil of 11 ablations (no lattice) and 20 ablations (with lattice) from n=3 biological replicates each. **e** sequential ablation: short junctional ablation followed by a longer 17µm ablation across the same junction. The latter one ablating the cortex connected to the ablated junction. **f** Quantification of vertex distance increase of sequential ablations. Line: mean +/-SD recoil of 7 sequential ablation from N=2 biological replicates. **g** Cortical laser ablation showing straightening of curved cell-cell boarders (yellow line) after linear ablation of the lattice.

### αII-spectrin enhances epidermal barrier formation

Although a differentiation hallmark, whether suprabasal cell flattening is sufficient to promote epidermal differentiation is still unclear (Simpson et al., 2011a; Luxenburg and Zaidel-Bar, 2019). We thus asked whether the inability to properly flatten upon depletion of αII-spectrin also changes epidermal differentiation. In E17.5 control embryos, staining for keratin (K)14 only marked the basal layer whereas in *Sptan1^KD^* epidermis also suprabasal cells were K14^+^ (Fig. 5a arrow; Supplementary Fig. 5a,b). In agreement, EdU incorporation assays showed that suprabasal but not basal proliferation was increased three-fold compared to control (Supplementary Fig. 5d, e). In contrast, the suprabasal marker K10 was not obviously altered, indicating at least a partial induction of differentiation (Fig. 5a; Supplementary Fig. 5a). Basal spindle orientation that can regulate cell fate (Williams et al., 2011) was also not altered (Supplementary Fig. 5c). Thus, in addition to shaping cell morphology, spectrin is critical for the initiation of basal cell differentiation during suprabasal translocation. We next asked whether the ability to induce differentiation is linked to changes in cell shape by analyzing K14⁺ basal and suprabasal cells in control and *Sptan1^KD^* epidermis. This analysis showed that especially suprabasal K14⁺ cells exhibited a defect in flattening whereas the increase in size upon SpectrinKD was less dependent on the differentiation status (Fig. 5b). Further, in *Sptan1^KD^* embryos, loricrin was detected predominantly at the cell periphery in the granular layer cells (Fig. 5a; Supplementary Fig. 5b), in contrast to control where loricrin^+^-granules filled the entire cytoplasm, suggesting impaired terminal differentiation and barrier formation.

**Figure 5.**
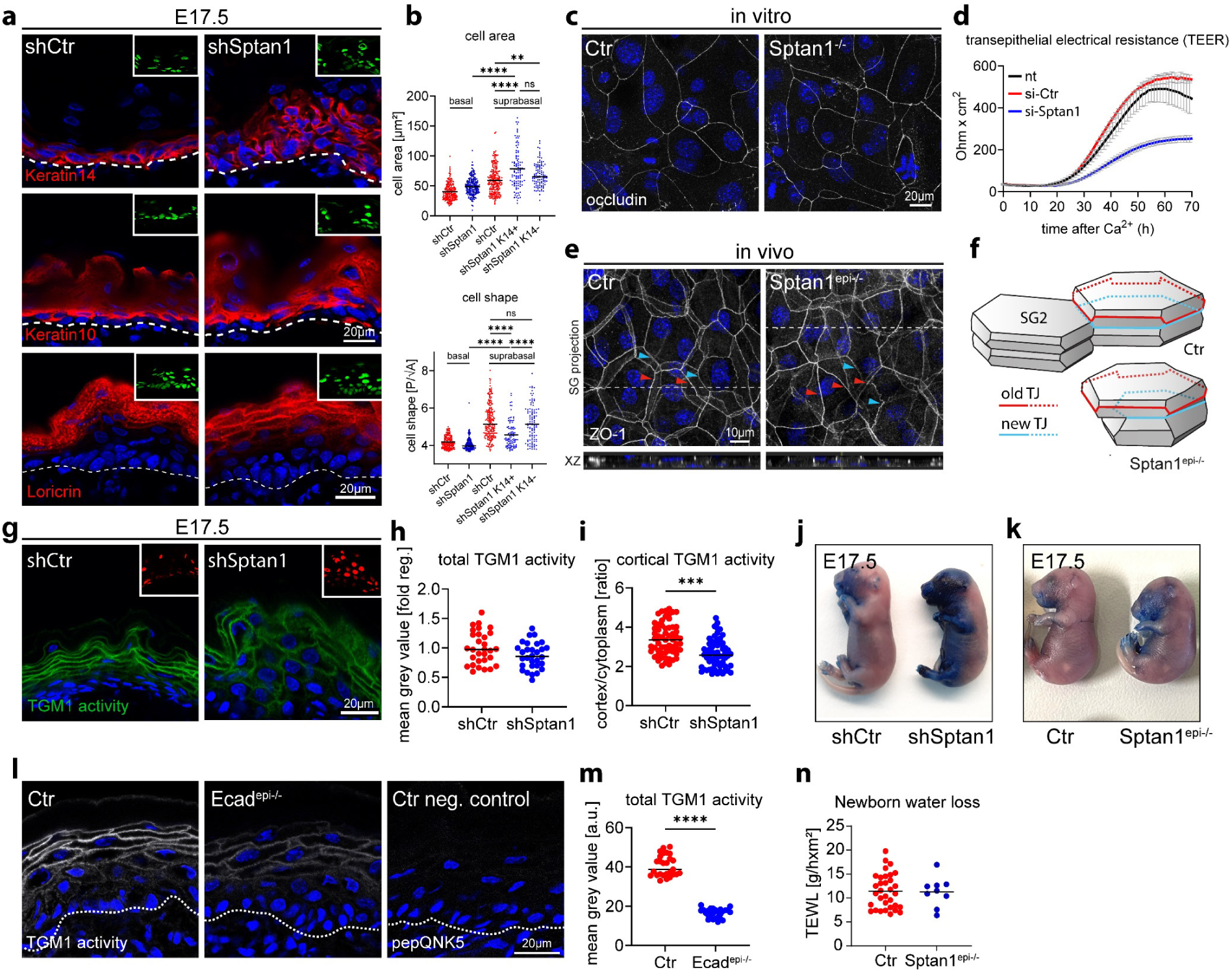
αII-spectrin regulates epidermal differentiation and barrier function. **a** Dorsal skin from *shCtr* and *shSptan1 0595* transduced E17.5 embryos immunolabeled for the basal layer marker K14, the differentiation marker K10 and the granular layer marker loricrin. Insets show the transduced cells (H2B−GFP+). **b** Quantification of cell area and shape of suprabasal *shSptan1* K14^+^ and K14^-^ cells in E17.5 embryos. Values of basal and suprabasal *shCtr* and basal *shSptan1* are equal to Fig. 1 Lines: Mean values; Dots: single cells pooled from 3 embryos with >100 cells for each condition. \*\*\*\**P* < 0.0001 with Kruskal–Wallis, Dunn’s multiple comparison test. **c** Immunofluorescence analysis for the TJ marker occludin in Ctr and Spectrin^-/-^ primary keratinocytes differentiated for 48h in high Ca^2+^. Representative example of 3 biological replicates each. **d** Transepithelial resistance (TER) measurements in Ctr and αII-spectrin knockdown keratinocytes after switching to high Ca^2+^. Line represent means over time of 3 biological replicates each. Representative experiment of n > 10 biological replicates. **e** Newborn epidermal whole-mount immunofluorescence analysis for the TJ marker ZO-1 revealing impaired alignment of the upper old (red arrowheads) and the lower new TJ rings (blue arrowheads) in the granular layer 2 (SG2). Maximum projection of the granular layer. **f** Illustration of cell shapes and TJ organization in the SG2 of Ctr and *Sptan1^epi-/-^* epidermis. **g** Dorsal skin sections from *shCtr* and *shSptan1 0595* transduced E17.5 embryos. Sections were processed for Transglutaminase1 (TGM1) activity assay. Upper Insets show the transduced cells (H2B−RFP+). **h** Quantification of TGM1 intensity from data shown in **g**. Data are the mean ± SD of 30 ROI from n=3 embryos per condition. Bars: mean normalized intensity; dots individual microscopy fields. \**P*=0.0471 by unpaired t-test. **i** Quantification of TGM1 activity cortical enrichment from the data shown in **g**. Mean ± SD from 60 individual cells from n=3 embryos per condition. Bars: TGM1 activity cortex/cytoplasm intensity ratio mean; dots: individual cells. \*\*\*\**P*<0.0001 by unpaired t-test. **j** Dye exclusion assay: *shCtr* and *shSptan1 0595* transduced E17.5 embryos were treated with toluidine blue dye to evaluate the skin barrier. **k** Dye exclusion assay: Ctr and *Sptan1^epi-/-^* E17.5 embryos were treated with toluidine blue dye to evaluate the skin barrier. **l** Dorsal skin section from Ctr and E-cadherin^epi-/-^ newborn mice. Sections were processed for Transglutaminase1 (TGM1) activity assay or negative Ctr (mutated TGM substrate, pepQNK5). **m** Quantification of TGM1 intensity from data shown in **l**. Data are the mean of 27 fields of view from n=3 newborn mice per condition. Bars: Mean intensity; dots individual microscopy fields. \*\*\*\**P*<0.0001 with Kolmogorov-Smirnov.

TJ formation is a key feature of the SG2 layer and necessary for epidermal barrier function (Kubo et al., 2009; Rübsam et al., 2017). To follow the formation of a functional TJ barrier and determine whether spectrin is required, we stratified primary keratinocytes and measured transepithelial electrical resistance (TEER). Although depleting spectrin did not obviously alter the recruitment of the TJ marker occludin to the apical cell-cell contacts (Fig. 5c), TEER remained low over time (Fig. 5d), thus indicating an inability to form a functional TJ barrier that could not be explained by lower cell numbers (Supplementary Fig. 5f). In the mouse epidermis, the tetrakaidecahedron-shaped SG2 cells that transition out of the SG2 layer not only have mature apical TJs but also form new TJs at basal tricellular contacts (Yokouchi et al., 2016b). In control the lower, nascent TJ ring is smaller but is aligned with and mirrors the shape of the upper, mature TJ ring (Fig. 5e, Ctr) due to their geometric stacking arrangement (Yokouchi et al., 2016b). This shape correlation and alignment was lost upon loss of αII-spectrin (Fig. 5e,f), reflecting a disrupted coordination between SG2 and SG3 cells as a result of altered cell shape which may explain dysfunctional TJs.

To further explore changes in terminal differentiation we assessed the activity of the cross-linker enzymes transglutaminases (TGMs) in the granular layer, necessary for terminal differentiation and proper formation of the outermost stratum corneum barrier (Eckert et al., 2005)(Simpson et al., 2011). Incubation with a fluorescent substrate peptide to localize TGM1 activity (Sugimura et al., 2008) showed high enrichment of this peptide at the cell cortex of control granular layer cells which was reduced by ∼30% reduction in *Sptan1^KD^* E17.5 embryos or 36% in *Sptan1^epi-/-^*newborns (Fig. 5g-I; Supplementary Fig. 5g, h). Similarly, *Ecad^epi-/-^* epidermis also showed decreased TGM activity (Fig. 5l,m) with no change in TGM1 protein levels upon epidermal loss of either αII-spectrin or E-cadherin (Supplementary Fig. 5i-k). Thus, E-cadherin-dependent cortical organization of actomyosin-spectrin networks directs TGM activation necessary for terminal differentiation (Supplementary Fig. 5k,l). Toluidine blue exclusion assays (Hardman et al., 1998) showed no dye penetration in control E17.5 embryos except for the most ventral side, indicating the formation of an intact barrier (Hardman et al., 1998). In contrast, E17.5 embryos from *Sptan1^KD^* and *Sptan1^epi-/-^* showed staining in the head and larger ventral parts of the embryo (Fig. 5j, k; Supplementary Fig. 5l) with ectopic staining still at the head region of E18.5 *Sptan1^KD^* embryos (Supplementary Fig. 5m) indicating a delay in stratum corneum barrier formation. However, transepidermal water loss was not obviously changed in *Sptan1^epi-/-^* newborns (Fig. 5n), suggesting that the developmental defect in barrier formation is a transient. Taken together, spectrin regulates cell shape and differentiation to enable functional TJ organization and cortical TGM activation downstream of E-cadherin necessary for proper barrier formation.

**Supplementary Figure 5.**
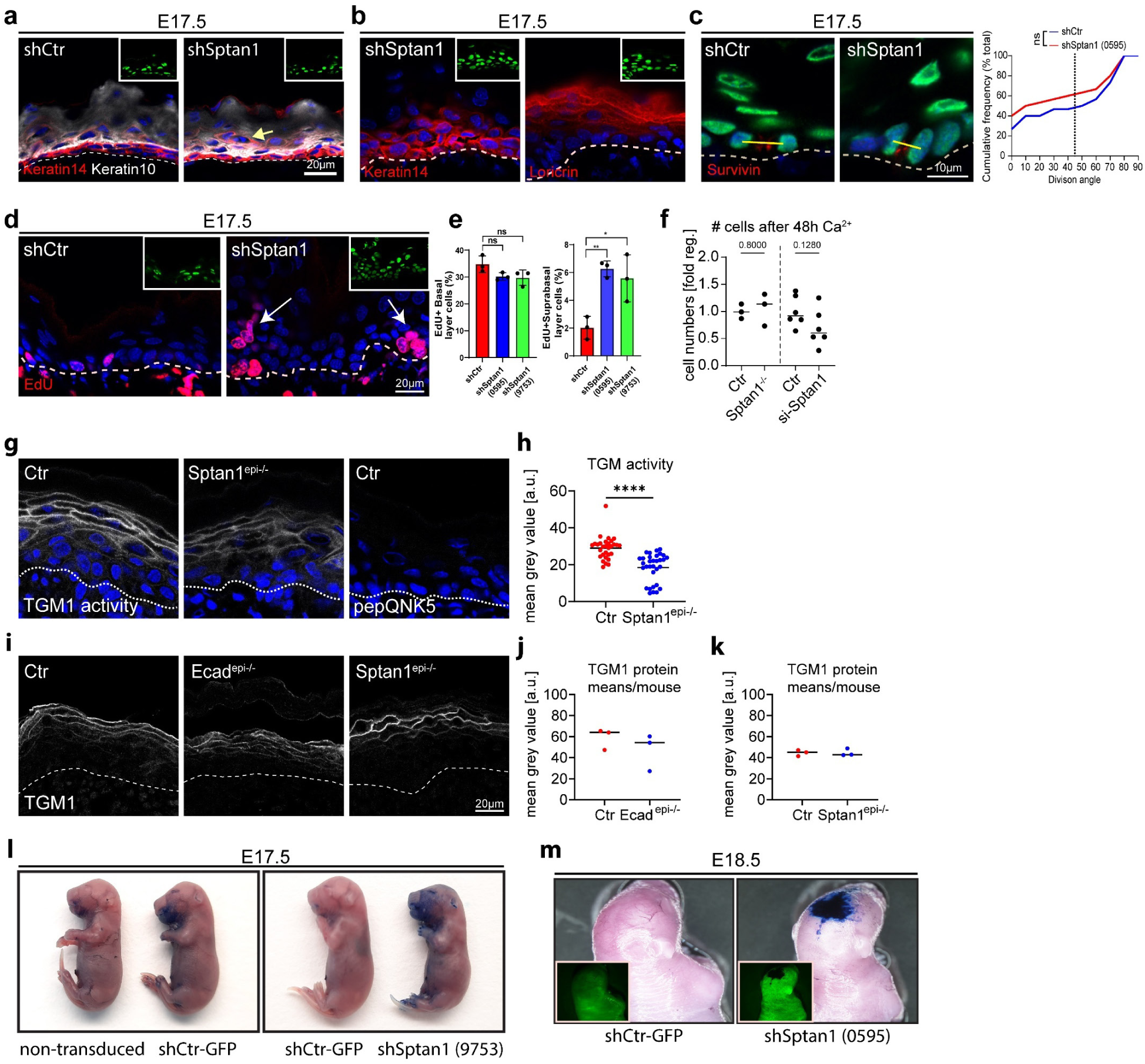
αII-spectrin regulates epidermal differentiation and barrier function. **a** Dorsal skin sections from *shSptan1 0595* transduced E17.5 embryos immunolabeled for the basal layer marker K14 and the suprabasal marker K10. **b** Dorsal skin sections from *shSptan1 9753* transduced E17.5 embryos immunolabeled for the basal layer marker K14 and the granular layer marker loricrin. Insets show the transduced cells (H2B−GFP+). **c** Dorsal skin sections from *shSptan1 0595* transduced E17.5 embryos immunolabeled for the cleavage furrow marker survivin. Yellow lines show representative axes of division. Graph: Quantification of spindle orientation plotted as a cumulative frequency distribution. NS: *P*=0.4 by Kolmogorov–Smirnov. **d** Dorsal skin sections from *shSptan1 0595* transduced E17.5 embryos immunolabeled for EdU. **e** Quantification of EdU+ basal and suprabasal layer cells from the data shown in **d**. Bars: mean ± SD from n=3 embryos per condition. Dots: average Edu^+^ basal and suprabasal layer cells from each embryo. NS: *P*=0.11 and NS: *P*=0.1097 for EdU+ basal layer cells. **P=0.002, *P=0.048 for EdU+ suprabasal layer cells by unpaired t-test. **f** Quantification of cell (nuclei) numbers from primary Ctr and *Sptan1^-/-^* or *Sptan1* siRNA treated keratinocytes differentiated for 48h in high Ca^2+^. Dots: Mean values from biological replicates. >360 cells counted for Ctr/*Sptan1^-/^*^-^ each and >20000 cells for siCtr/*siSptan1* each. **g** Dorsal skin section from Ctr and *Sptan1^epi-/-^*newborn mice. Sections were processed for Transglutaminase1 (TGM1) activity assay or negative Ctr (mutated TGM substrate, pepQNK5). **h** Quantification of TGM1 intensity from data shown in **g**. Data are the mean of 30 fields of view from n=3 newborn mice per condition. Bars: Mean intensity; dots individual microscopy fields. \*\*\*\**P*<0.0001 with Kolmogorov-Smirnov. **i** Dorsal skin sections from Ctr, *Ecad^epi-/-^* and *Sptan1^epi-/-^* newborn mice immunolabeled for total TGM1 protein. **j, k** Quantification of TGM1 intensity in Ctr, *Ecad^epi-/-^*and *Sptan1^epi-/-^*. Lines: Mean values/biological replicate. Non-significant with Mann Whitney. **l** Dye exclusion assay: *shCtr* and *shSptan1 9753* transduced E17.5 embryos were treated with toluidine blue dye to evaluate the skin barrier. **m** Dye exclusion assay: *shCtr* and *shSptan1 0595* transduced E18.5 embryos were treated with toluidine blue.

### αII-spectrin is required for the activity of the EGFR-TRPV3-TGM pathway

We previously showed that E-cadherin controls epidermal barrier formation by regulating the epidermal growth factor receptor (EGFR) (Rübsam et al., 2017). Furthermore, EGFR activation promotes the opening of the calcium channel TRPV3 necessary for TGM1 activation and subsequent terminal epidermal differentiation (Cheng et al., 2010). As spectrin functions as a cortical platform that organizes membrane domains and membrane protein activities (Machnicka et al., 2014), we hypothesized that αII-spectrin promotes terminal differentiation through the EGFR-TRPV3 pathway. Immunofluorescence analysis showed that EGFR was expressed in all epidermal layers in both control and *Sptan1*^KD^ E17.5 embryos, with, as expected, most intense staining in the basal layer (Suppl. Fig. 6a). Whereas low levels of phosphorylated EGFR (pEGFR) were detected in control basal and spinous layers, the most intense pEGFR staining was seen at the cortex of granular layer cells. In contrast, many *Sptan1*^KD^ granular layer cells showed diffuse or no pEGFR staining (30% decrease in pEGFR intensity) with abnormally strong cortical staining now detected in spinous layer cells (4-fold increase)(Fig. 6a-c).

**Figure 6.**
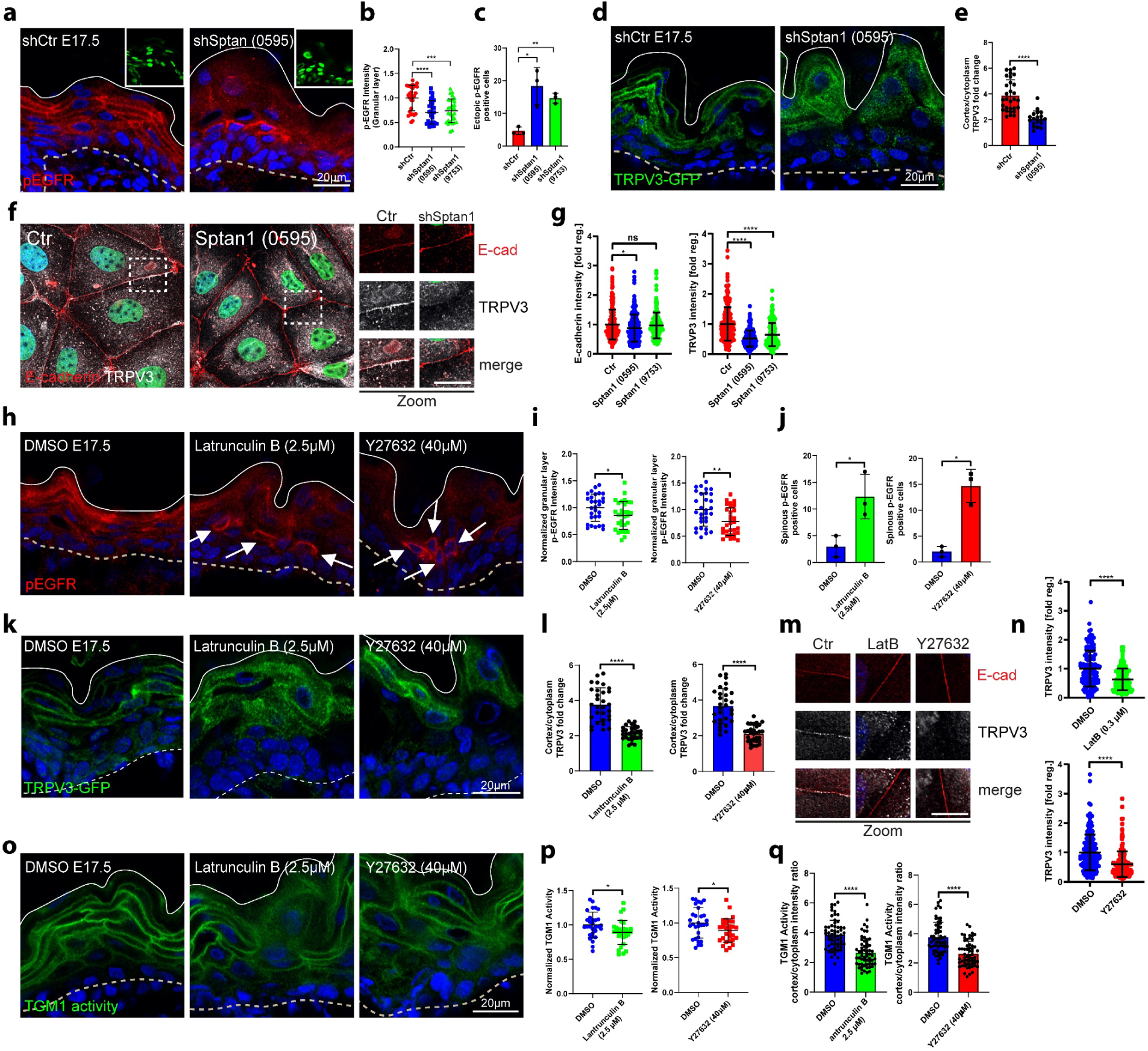
αII-spectrin-actomyosin networks regulate the EGFR-TRPV3-TGM pathway. **a** Dorsal skin sections from *shCtr*, and *shSptan1 0595* transduced E17.5 embryos immunolabeled for pEGFR. Upper Insets show transduced cells (H2B−GFP+). **b** Quantification of pEGFR granular layer intensity from data shown in **a**. Mean ± SD of 30 ROI (Region of interest) from n=3 embryos per condition. Horizontal bars: mean normalized intensity, dots: individual microscopy fields. \*\*\*\**P*<0.0001, \*\*\**P*=0.0001 by unpaired t-test. **c** Quantification of ectopic pEGFR-positive cells from data shown in **a**. Horizontal bars: mean ± SD from n=3 embryos per condition, dots: average ectopic pEGFR-positive cells per embryo. \**P*=0.0482, \*\**P*=0.0011 by unpaired t-test. **d** Sections of dorsal skin from *shCtr;* TRPV3-GFP and *shSptan1 0595*; TRPV3-GFP transduced E17.5 embryos. **e** Quantification of TRPV3-GFP cortical enrichment from the data shown in **d**. Mean ± SD from 30 individual cells from n=3 embryos per condition. Bars: TRPV3-GFP cortex/cytoplasm ratio, dots: individual cells. \*\*\*\**P*<0.0001 by unpaired t-test. **f** *ShCtr* and *shSptan1-0595* transduced primary mouse keratinocytes (H2B-GFP+) cultured in high-calcium (1.5 mM) medium and immunolabelled for E-cadherin and TRPV3. Insets (right) show magnifications of the boxed areas. **g** Quantification of E-cadherin and TRPV3 intensity. Mean ± SD from ∼200 mature junctions from n=3 experiment per condition. Bars: mean normalized intensity, dots: mature junctions. *P<0.05, ****P<0.0001 by unpaired t-test. **h** Dorsal skin sections from E17.5 wild-type embryos treated with DMSO, latrunculin, or Y27632 immunolabeled for p-EGFR. **i** Quantification of pEGFR granular layer intensity from data shown in **h**. Mean ± SD of 30 ROI from n=3 embryos per condition. Bars: mean normalized intensity, dots: individual microscopy fields. \**P*=0.0345 (Latrunculin B), \*\**P*=0.0035 (Y27632) unpaired t-test. **j** Quantification of ectopic pEGFR-positive cells from data shown in **h**. Bars: mean ± SD from n=3 embryos per condition. Dots: average ectopic pEGFR-positive cells from each embryo. \**P*=0.0422 (Latrunculin B), \**P*=0.0143 (Y27632) by unpaired t-test. **k** Dorsal skin section from *shCtr;* TRPV3-GFP transduced E17.5 embryos treated with DMSO, latrunculin or Y27632. Insets show the magnification of the boxed area from each epidermal layer. **l** Quantification of TRPV3-GFP cortical enrichment from the data shown in **k**. Data are the mean ± SD from 30 individual cells from n=3 embryos per condition. Horizontal bars represent the TRPV3-GFP cortex/cytoplasm intensity ration mean and circles represent individual cells. \*\*\*\**P* <0.0001 by unpaired t-test. **m** Primary mouse keratinocytes cultured in high-calcium (1.5 mM) medium treated with DMSO, latrunculin or Y27632 and immunolabelled for E-cadherin and TRPV3. **n** Quantification of TRPV3 intensity from data shown in C. Mean ± SD from ∼150 mature junctions from n=3 experiment per condition. Bars: Mean normalized intensity, dots: mature junctions. ****P <0.0001 by unpaired t-test. **o** Dorsal skin sections from E17.5 wild-type embryos treated with DMSO, latrunculin or Y27632 and processed for Transglutaminase activity assay. **p** Quantification of crosslinked TGM substrate intensity. Mean ± SD of 30 ROIs from n=3 embryos per condition. Bars: Mean normalized intensity, dots: individual microscopy fields. \**P*=0.0205 (Latrunculin B), \**P*=0.0345 (Y27632) by unpaired t-test. **m** Quantification cortically enriched crosslinked TGM substrate. Mean ± SD from 60 individual cells from n=3 embryos per condition. Bars: means of cortex/cytoplasm ratio, dots: individual cells. \*\*\*\**P*<0.0001 by unpaired t-test. Nuclei were stained with DAPI; dotted lines indicate the dermal-epidermal border.

To determine whether αII-spectrin controls TRPV3 localization, we infected E9 embryos with lentivirus encoding TRPV3-GFP (Xiao et al., 2008). In control E17.5 epidermis, TRPV3^GFP^ was diffuse in the basal and spinous layers and became cortical in the granular layer (Fig. 6d) where it co-localized with pEGFR (Supplementary Fig. 6b). In contrast, in *Sptan1*^KD^ embryos, TRPV3^GFP^ remained more diffuse in the granular layer (50% reduction in cortical enrichment, Fig. 6d; Supplementary Fig. 6c). Furthermore, in control primary keratinocytes cells, endogenous TRPV3 colocalized with E-cadherin at the cell membrane, which was significantly reduced in *Sptan1*^KD^ cells (Fig. 6f,g). Thus, spectrin is essential for cortical activation of EGFR and for cortical localization of TRPV3.

### αII-spectrin-actomyosin networks regulate the EGFR-TRPV3-TGM pathway

Given that spectrin directs the spatial organization and mechanics of the cortical cytoskeleton, we asked whether αII-spectrin regulates the EGFR-TRPV3-TGM axis through actomyosin. To this end, we treated *shCtr* E17.5 embryos with latrunculin, and Y27632 to decrease F-actin levels and myosin II motor activity, respectively. Similar to depletion of αII-spectrin, latrunculin or Y27632 treatment decreased pEGFR levels in the granular layer (15 and 23%, respectively), and increased (4- or 7-fold) spinous cortical pEGFR levels relative to control, DMSO-treated, embryos (Fig. 6h-j). Importantly, latrunculin or Y27632 also decreased the enrichment of TRPV3^GFP^ in vivo (50%, Fig. 6k,l) and the recruitment of endogenous TRPV3 to intercellular contacts in cultured keratinocytes (Fig. 6m,n; Supplementary Fig. 6d, e). Moreover, these treatments further decreased cortical enrichment of the TGM1 substrate peptide by ∼40% in E17.5 embryos (Fig. 6o-q). Together, the results demonstrate that E-cadherin-dependent recruitment of spectrin controls the organization and mechanics of the cortical cytoskeleton in suprabasal layers, which, in turn, determines cell shape and EGFR and TRPV3 activity at the membrane necessary to activate TGM in the granular layer required for epidermal barrier formation.

**Supplementary Figure 6.**
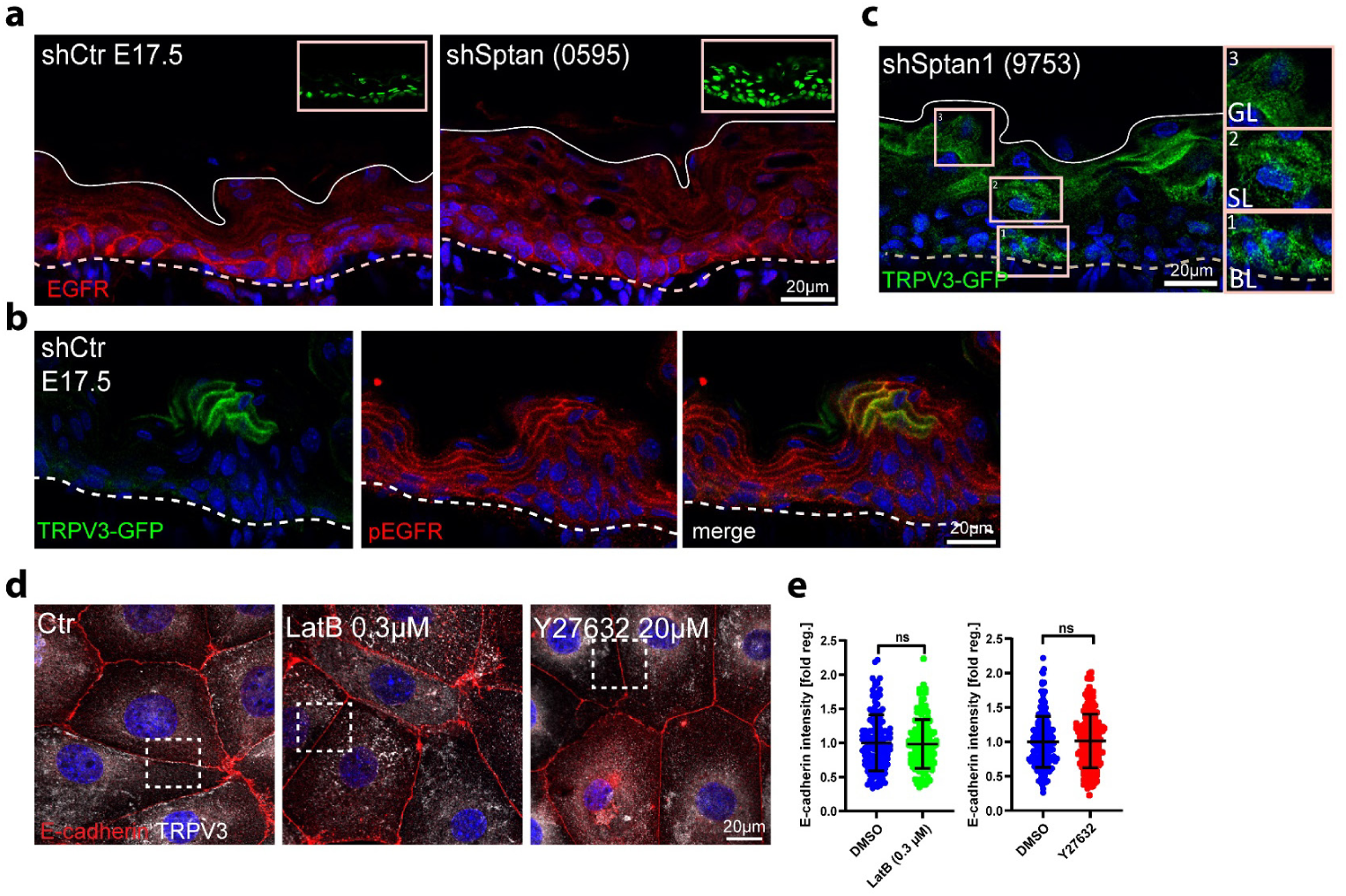
αII-spectrin-actomyosin networks regulate the EGFR-TRPV3-TGM pathway. **a** Dorsal skin sections from *shCtr* and *shSptan1 0595* transduced E17.5 embryos immunolabeled for EGFR. Upper Insets show the transduced cells (H2B−GFP+). **b** Dorsal skin sections from *shCtr;* TRPV3-GFP transduced E17.5 embryos immunolabeled for pEGFR. **c** Dorsal skin sections from *shSptan1 9753*; TRPV3-GFP transduced E17.5 embryos. Insets show the magnification of the boxed area from each epidermal layer. **d** Primary mouse keratinocytes cultured in high-calcium (1.5 mM) medium treated with DMSO, latrunculin or Y27632 and immunolabelled for E-cadherin and TRPV3. Overviews corresponding to Fig. 6m. Boxes indicate the location of magnified area. **e** Quantification of E-cadherin intensity. Mean ± SD from ∼150 mature junctions from n=3 experiment per condition. Bars: Mean normalized intensity, dots: mature junctions. Nuclei were stained with DAPI.

### EGFR activity regulates cortical TRPV3 localization

To further explore the relationship between spectrin-actomyosin cortical organization, TRPV3 localization and localized EGFR activity, we inhibited EGFR using gefitinib in either E17.5 TRPV3^GFP^-expressing embryos or in cultured keratinocytes. Compared to DMSO control, gefitinib treatment significantly reduced cortical enrichment of both αII-spectrin and TRPV3 at intercellular contacts (Fig. 7d-g). Interestingly, activation of EGFR by TGFα treatment also significantly decreased αII-spectrin and TRPV3 localization at intercellular junctions (Fig. 7d-g).

**Figure 7.**
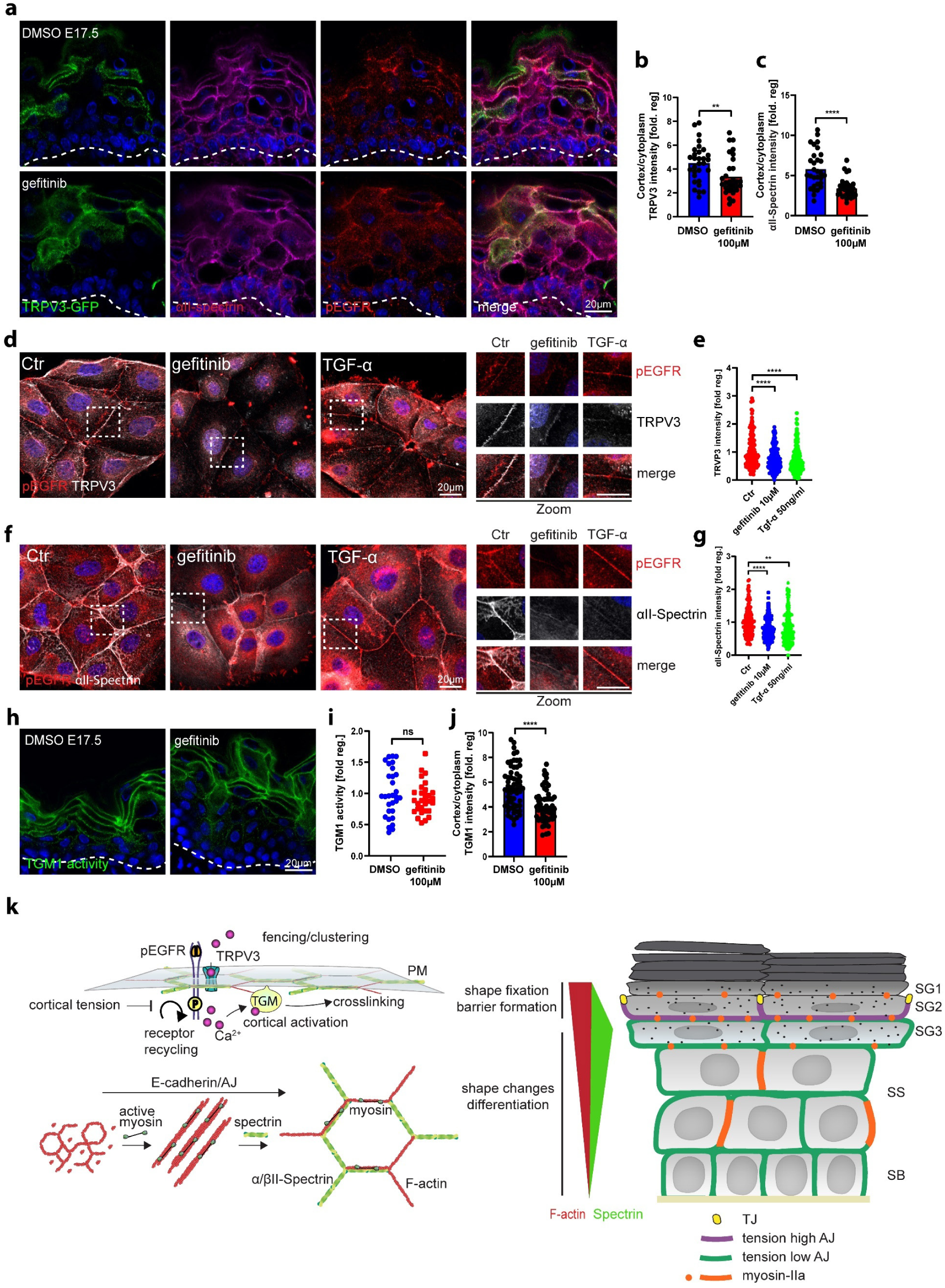
EGFR activity regulates cortical TRPV3 localization. **a** Dorsal skin sections from shCtr; TRPV3-GFP transduced E17.5 embryos treated with DMSO or gefitinib immunolabeled for αII-spectrin and pEGFR. **b** and **c** Quantification of TRPV3-GFP and αII-spectrin cortical enrichment from the data shown in **a**. Mean ± SD from 30 individual cells from n=3 embryos per condition. Bars: TRPV3-GFP cortex/cytoplasm ratio; dots: individual cells. \*\**P*=0.0024 for TRPV3-GFP and **** *P*>0.0001 for αII-spectrin by unpaired t-test. **d** and **f** Primary mouse keratinocytes cultured in high-calcium (1.5 mM) medium treated with DMSO, gefitinib or TGF-α and immunolabelled for p-EGFR, TRPV3 and αII-spectrin. Boxes indicate the location of magnified area. **e** and **g** Quantification of TRPV3 and SPTAN1 intensity from the data shown in **d** and **f**. Mean ± SD from ∼200 mature junctions from n=3 experiment per condition. Bars: Mean normalized intensity; dots: individual junctions. Nuclei were stained with DAPI. **** *P*>0.0001 for TRPV3 intensity. **** *P*>0.0001 and \*\**P*=0.0013 for αII-spectrin intensity. **h** Dorsal skin sections from E17.5 wild-type embryos treated with DMSO and gefitinib. Sections were processed for Transglutaminase1 (TGM1) activity assay. **i** Quantification of crosslinked TGM substrate intensity. Mean ± SD of 30 ROIs from n=3 embryos per condition. Bars: Mean normalized intensity; dots: individual microscopy fields. NS: *P*=0.3473 by unpaired t-test. **j** Quantification of cortical enrichment of crosslinked TGM substrate. Mean ± SD from 60 individual cells from n=3 embryos per condition. Bars: means of cortex/cytoplasm ratio, dots: individual cells. \*\*\*\**P*<0.0001 by unpaired t-test. Nuclei were stained with DAPI; dashed lines indicate the dermal-epidermal border. **k** Model - High-tension actomyosin-spectrin cortices regulate transglutaminase activity. Illustration of junction and cytoskeleton distribution across epidermal layers with spectrin most enriched in SG3 and F-actin most enriched in SG1. Lower left: illustration of spectrin and myosin dependent organization of cortical F-actin. Upper left: Working model how lattice organization and myosin tension regulate EGFR/TRPV3 signaling complexes resulting in Ca^2+^ influx and cortical transglutaminase activation.

Finally, we asked whether EGFR activity is sufficient to regulate TGM1 activity in the epidermis. As in *Sptan1^KD^* embryos (Fig. 5g-i), cortical enrichment of the TGM1 substrate peptide was markedly decreased upon gefitinib treatment, indicating reduced cortical TGM1 activity (Fig. 7h-j). Taken together, these and previous findings (Rübsam et al., 2017; Cheng et al., 2010) highlight a critical interdependency between the organization and tensile state of the actomyosin-spectrin cortex that not only determines cell shape but also spatially confines EGFR/TRPV3 activity at the cell membrane only in the granular layer to induce terminal differentiation and establish a functional epidermis.

## Discussion

In 1858, physician Rudolf Virchow established a correlation between cell shape and function, describing how abnormalities in cell shape can lead to disease development (Virchow, 1871). While the regulation of cell shape is a well-studied process dependent on adhesion and actomyosin, the mechanisms by which these processes also regulate differentiation and function require further elucidation. The layered structure of the epidermis provides a unique example of the correlation between cell position, shape, and function, i.e., differentiation and barrier function (Peskoller et al., 2022; Luxenburg and Zaidel-Bar, 2019). This study demonstrates that E-cadherin-dependent integration of αII-spectrin into submembraneous actomyosin networks is crucial for the differential organization and tension state of the cortex to coordinate cell shape and differentiation state in each layer essential to form a functional skin barrier.

Previously, we demonstrated that E-cadherin, a master regulator of cell-cell adhesion, is a key regulator of actomyosin-dependent tension to spatially control the localization and activation of EGFR, essential for TJ barrier function in the epidermis (Rübsam et al., 2017). However, how E-cadherin controls polarized localization of actomyosin tension and EGFR activity across layers remained unclear. Our new data now show that differential gradients of F-actin, myosin II and spectrin drive a layer-specific organization of honeycomb-shaped spectrin-actomyosin networks necessary to dissipate myosin-dependent tension. Further, spectrin’s elastic properties likely allow the cortex to not only accommodate the change in cell shape when moving into a new layer, but also to absorb forces experienced during this movement. Finally, this layer-specific organization of spectrin-actomyosin cortical network regulates the localization and activity of EGFR/TRPV3 Ca^2+^ channel complexes necessary to activate TGM only in the granular layer to promote terminal differentiation and epidermal barrier function. Our data thus link cell adhesion with position-dependent cytoskeletal configuration to direct spatial control of cell shape, cell fate and barrier function.

Previous studies on cell shape-dependent control of differentiation in the epidermis focused on early differentiation of basal progenitor cells and involved regulation of spindle orientation as well as compression-induced delamination (Le et al., 2016; Luxenburg et al., 2011; Miroshnikova et al., 2018). In contrast, other studies in which different actin-binding proteins, such as cofilin, WDR1, thymosin β4 and others, were deleted in the epidermis did not show any obvious changes in global differentiation markers (Luxenburg et al., 2015; Padmanabhan et al., 2020; Mahly et al., 2022;). Similarly, K10 was not obviously altered upon loss of αII-spectrin even if the basal marker K14 was upregulated. Our data now show that changes in cortical spectrin-actomyosin organization not only control basal cell shape and initial differentiation but drive suprabasal cell shape transitions to regulate terminal differentiation, thus coordinating cell shape and fate across the tissue.

How spectrin regulates early differentiation when transitioning in the spinous layer is less clear. One potential candidate is Yap, based on previous studies that implicated spectrin and actomyosin tension in controlling YAP signaling in other epithelia (Deng et al., 2015; Fletcher et al., 2015), However, YAP nuclear localization was unchanged in *Sptan1*-depleted epidermis, suggesting that spectrin controls differentiation through a YAP-independent mechanism.

In the epidermis, mechanical regulation of EGFR contributes to barrier function by promoting terminal differentiation through TRPV3 and TGM1 (Cheng et al., 2010) and regulating TJs by reinforcing high tension states in the SG2 layer (Rübsam et al., 2017) but whether these two functions are facets of the same mechanical regulation or regulated through distinct mechanisms remains unclear. Previous studies have shown that loss of myosin IIa/b in the epidermis impaired TJ barrier function but did not obviously affect stratum corneum barrier function (Sumigray et al., 2012), whereas loss of the Arp2/3 complex that enhances branched actin network assembly affects both barriers (Zhou et al., 2013). These observations further demonstrate the profound complexity of the actomyosin cytoskeleton and the importance of its fine-tuning for the execution of distinct biological processes.

Our previous and current data indicate that layer-specific organization of E-cadherin-directed spectrin-actomyosin lattices also controls the activity and localization of EGFR, with active EGFR being internalized in lower layers whereas in the granular layer active EGFR is retained at the membrane. Either loss of spectrin or reducing actomyosin tension reverses this localization of active EGFR, with membrane localization now being confined to lower layers. The organization of the spectrin-actomyosin cortex can inhibit diffusion of plasma membrane receptors to induce molecular crowding, known as fencing (Fujiwara et al., 2016), as well as organize membrane trafficking events such as endocytosis (Ghisleni et al., 2020). We thus propose a model in which the spectrin-actomyosin organization in the granular layers limits mobility of EGFR and TRPV3, while the high tension state of this network also inhibits internalization of activated EGFR, thus bringing these proteins in close proximity to activate TRPV3 and, subsequently, TGM1 (Fig.7K). In agreement, both ligand binding and actin-dependent membrane organization regulates EGFR distribution at the plasma membrane (Sankaran et al., 2021; Bag et al., 2015). A potential alternative model involves a mechano-sensitive unfolding of spectrin (Leterrier and Pullarkat, 2022; Renn et al., 2019) or other, unknown players within the granular layer cortex that then interact with and cage the EGFR/TRPV3 to the cortex. Thus far, however, there is no evidence for tension-dependent binding partners that may explain EGFR or TRPV3 retention.

In conclusion, our data provide evidence for a model in which E-cadherin-dependent integration of αII-spectrin into the actomyosin network spatially coordinates cortical organization and mechanics to determine cell shape and control localization of pEGFR-TRPV3 complexes at the plasma membrane resulting in TGM-dependent crosslinking only in the granular-layer, thereby fixing cell shapes and promoting stratum corneum barrier function.

## Materials and Methods

### Mice

To generate epidermal E-cadherin knockout (*Ecad^epi−/−^)* BL/6N mice, female *Cdh1^flox/flox^* mice were crossed to male *Ecad^fl/wt^*;K14-Cre mice. Transgenic *Ecad^flox/flox^* mice and K14-Cre mice and their genotyping were described previously (Hafner et al., 2004; Boussadia et al., 2002). Hsd:ICR (CD1) mice (Envigo) were used for all in vivo knockdown experiments. To generate epidermal αII-Spectrin knockout (*Sptan1^epi-/-^*) mice, *Sptan1^flox^* mice were ordered from The Jackson Laboratory (Strain ID-B6;129S-*Sptan1^tm1.1Mnr^*/J, Strain #033392). Female *Sptan1^flox/flox^* mice were crossed to male *Sptan1^fl/wt^*;K14-Cre mice. *Sptan1^flox^* mice were described previously (Huang et al., 2017). To generate epidermal confetti BL/6N mice, female R26R-Confetti^tg/tg^ mice were crossed with male K14-Cre mice for stochastic activation and recombination of the of the brainbow2.1 cassette in the epidermis resulting in four possible outcomes: Expression of nuclear GFP, membrane associated CFP, cytoplasmic YFP or cytoplasmic RFP.

### Quantification of 3D epidermal cell shape parameter using epidermal confetti

Epidermal confetti newborn mice were sacrificed and fixed in 4% PFA over night at 4°C. Backskin was then taken and mounted in Mowiol. As a basis for rendering the cytoplasmic YFP signal of the brainbow2.1 was used due to homogenous intracellular expression and across epidermal layers and good signal/noise ratio. Confocal stacks were acquired using a Leica TCS SP8 (PlanApo 63x, 1.4 NA CS2) in Leica lightning mode, Pinnhole 0.97AU, voxel size 0.046×0.046×0.259 followed by the built-in lightning deconvolution. Imaris was then used for surface rendering using a surface detail of 0.2µm and smoothing. Cell volume, surface area and sphericity were extracted from the rendered cells in Imaris. Cell position was determined from XZ projections of the same stack using the measure tool in Fiji. Cell position was determined relative to the stratum corneum by measuring the distance of each cell to the upper side of the stratum corneum. The stratum corneum boarder was readily detectable due to preservation of the fluorescent signal and high cellularity.

### Isolation and culture of primary keratinocytes

Primary keratinocytes were isolated and cultured as described before (Rübsam et al., 2017): newborn mice were decapitated and incubated in 50% betaisodona/PBS for 30min at 4°C, 1 min PBS, 1 min 70% EtOH, 1min PBS and 1 min antibiotic/antimycotic solution. Skin was incubated in 2ml dispase (5 mg ml^-1^ in culture medium) solution. After incubation over night at 4 °C, skin was transferred onto 500 µl FAD medium on a 6 cm dish and epidermis was separated from the dermis as a sheet. Epidermis was transferred dermal side down onto 500 µl of TrypLE (Thermo Fisher Scientific) and incubated for 20 min at RT. Keratinocytes were washed out of the epidermal sheet using 3 ml of 10% FCS/PBS. After centrifugation keratinocytes were resuspended in FAD medium and seeded onto collagen type-1 (0.04 mg ml^-1^) (Biochrom, L7213) coated cell culture plates. Primary murine keratinocytes were kept at 32 °C and 5% CO_2_.

Primary keratinocytes were cultured in DMEM/HAM’s F12 (FAD) medium with low Ca^2+^ (50 μM) (Biochrom) supplemented with 10% FCS (chelated), penicillin (100 U ml^-1^), streptomycin (100 μg ml^-1^, Biochrom A2212), adenine (1.8×10^−4^ M, SIGMA A3159), L-glutamine (2 mM, Biochrom K0282), hydrocortisone (0.5 μg ml^-1^, Sigma H4001), EGF (10 ng ml^-1^, Sigma E9644), cholera enterotoxin (0.1 nM, Sigma C-8052), insulin (5 μg ml^-1^, Sigma I1882), and ascorbic acid (0.05 mg ml^-1^, Sigma A4034).

### Transfection

Overexpression: Keratinocytes were transfected at 100% confluency with Viromer®Red. 1.5 µg DNA were diluted in 100 µl Buffer E, added to 1.25 µl Viromer®RED and incubated for 15 min at RT (Rübsam et al., 2017). 33 µl transfection mix were used per well with 0.5 ml FAD medium (24 well plate). Knockdown: Keratinocytes were transfected at 100% confluency with Viromer®BLUE (lipocalyx; Halle Germany). Transfection mix was prepared according the manufacturers protocol. 100 µl transfection mix was used per well with 1 ml FAD medium (12 well plate, 5 nM siRNA f.c.). siRNA (siPOOLs) against αII-Spectrin were obtained from (siPOOLs from siTOOLs, Planegg, Germany).

### Lentiviruses

Lentiviruses were produced as previously described (Beronja et al., 2010; Soffer et al., 2022). Briefly, lentiviral plasmids were generated by cloning oligonucleotides into pLKO.1-TRC (gift from David Root, Broad Institute, Cambridge, MA, USA; Addgene plasmid #10878), lentivirus-GFP or lentivirus-RFP (gift from Elaine Fuchs, Rockefeller University, New York, NY, USA; Addgene plasmid #25999) by digestion with EcoRI and AgeI, as described in the Genetic Perturbation Platform (GPP) website (http://portals.broadinstitute.org/gpp/public/resources/protocols). shRNA sequences were obtained from GPP (https://portals.broadinstitute.org/gpp/public/): *Sptan1(0595)* construct #TRCN0000090595, target sequence 5’-GCCACCGATGAAGCTTATAAA −3’. *Sptan1(9753)* construct #TRCN0000089753, target sequence 5’-CCAGTCACAATCACCAATGTT −3’. Ank3(6780) construct #TRCN0000236780, target sequence 5’-TGCCGTGGTTTCCCGGATTAA −3’. The TRPV3-GFP construct was a gift from Michael X. Zhu (UTHelath Houston).

### In-utero lentivirus injection

Lentiviruses were injected into gestating mice as previously described (Beronja et al., 2010). Briefly, females at E8.5 were anesthetized with isoflurane, injected with pain killer, Rheumocam Veterinary 5 mg/ml according to the manufacturer instructions (Chanelle Pharma, Irland) and each embryo (up to 6 per litter) was injected with 0.4 to 1 μl of approximately 2 × 10^9^ colony-forming units (CFUs) of the appropriate lentivirus. Controls were both uninfected littermates of *shSptan1*-*0595*/*9753*;H2B-GFP/*shSptan1*-*0595*;H2B-RFP/*shAnk3-6780*;H2B-GFP lentivirus-injected embryos and *shScr*;H2B-GFP lentivirus-injected embryos. For *shSptan1-0595/9753; TRPV3-GFP* lentivirus-injected embryos *shScr; TRPV3-GFP* were controls. In utero lentivirus injection infects approximately 60% to 70% of the dorsal skin epidermis (Beronja et al., 2010).

### Immunofluorescence of keratinocytes in vitro

Immunofluorescence stainings of keratinocytes were performed as described before (Sahu et al., 2020a): cells were seeded on collagen coated glass cover slips in a 24 well plate and switched to high Ca^2+^ medium as indicated in the results section. Cells were fixed using 4% PFA for 10 minutes at RT, washed three times for 5 minutes using PBS, permeabilized using 0.5% TritonX100/PBS and blocked using 5% NGS/1% BSA/PBS for 1 hour at room temperature. Primary antibodies were diluted as indicated in the antibody section in Background Reducing Antibody Diluent Solution (ADS)(DAKO). Cover slips were placed growth surface down onto a 50 ml drop of staining solution on parafilm in a humidified chamber and incubated overnight at 4 °C. Cover slips were washed again with PBS three times for 10 minutes. Secondary antibodies and DAPI (40,6-diamidin-2-phenylindol, Sigma) were diluted 1:500 in ADS and cover slips were incubated for 1 hour at RT. Secondary antibodies were washed off via three wash steps using PBS for 10 minutes. Cover slips were mounted using Mowiol (Calbiochem).

### Immunofluorescence on tissue sections

Embryos or newborn mice were embedded in OCT (Scigen), frozen, sectioned at 10 μM and fixed in 4% formaldehyde for 10 min. Sections were then blocked with 0.1% Triton X-100, 1% bovine serum albumin, 5% normal donkey serum in phosphate-buffered saline, or in MOM Basic kit reagent (Vector Laboratories). Sections were incubated with primary antibodies overnight at 4°C and with secondary antibodies for 1 h at room temperature.

### Antibodies and inhibitors

**Ankyrin3** (IF 1:500, WB 1:1000, Proteintech #27766-1-AP); **E-cadherin** (IF 1:200, BD Transduction Laboratories #610182, clone number 36 or IF 1:500, Cell Signaling #3195); **EGFR** (IF 1:500, Abcam #ab52894); pEGFR (Y1068)(IF 1:500, Abcam #ab40815); **GAPDH** (WB 1:10000, Ambion #AM4300 or WB 1:1000, Cell Signaling #5174); **GFP** (1:3000, Abcam #ab13970); **Keratin10** (IF 1:1000, BioLegend #PRB-159P); **Keratin14** (IF 1:2000, Covance #PRB 155P); **Loricrin** (1:1000, BioLegend #Poly19051); **Myosin heavy chain IIa** (IF 1:500, Biolegend PRB-440); **phospho-Myosin Light Chain 2** (Thr18/Ser19) (IF 1:100, Cell Signaling #3674); **occludin** (IF 1:400, Invitrogen #33-1500); **TRPV3** (IF 1:1000, Abcam #b94582); **αII-Spectrin** (IF 1:500, WB 1:1000, Abcam ab11755); **TGM1** (IF 1:500, proteintech #12912-3-AP); **α-catenin** (IF 1:2000, Sigma #C2081); **β-catenin** (IF 1:1000, Abcam #ab32572); **YAP** (IF 1:500, Cell Signaling #14074). **Phalloidin** was used to stain F-actin (IF 1:500, Sigma #P1951, TRITC conjugated). Secondary antibodies were species-specific antibodies conjugated with either AlexaFluor 488, 594 or 647, used at a dilution of 1:500 for immunofluorescence (Molecular Probes, Life Technologies), or with horseradish peroxidase antibodies used at 1:5000 for immunoblotting (Bio-Rad Laboratories). Inhibitors used in this study if not indicated otherwise: Blebbistatin myosin inhibitor, 5 μM (Sigma #B0560); Latrunculin B, 0.1 μM (Sigma L5288).

### Preparation of epidermal whole mounts

Epidermal whole mounts were prepared as has been described previously (Rübsam et al., 2017). Backskin was prepared from newborn mice and subcutaneous fat was removed with curved tweezers. The epidermis was mechanically, carefully peeled off from the dermis with ultrafine curved tweezers thereby separating the basal from the suprabasal layers. During the whole procedure the basal side of the sheet was kept floating on PBS2+ (PBS supplemented with 0.5mM MgCl2 and 0.1mM CaCl2). Subsequently the epidermal sheet was fixed floating on 4% PFA on ice for 10 min, washed on PBS for 5 min and permeabilized with 0.5% TritonX100/PBS for 1 h at room temperature (RT). The permeabilized sheet was washed for 5 min on PBS and blocked with 10% FCS/PBS for 30 min/RT. For staining epidermal sheet were cut into ca. 5 × 5mm pieces. The SC of the epidermis is prone unspecific binding of ABs. Additionally it cannot be permeabilized to allow AB permeation. Thus, all following steps were performed incubating the sheet from the basal side leaving the SC side dry. Primary ABs were diluted either in AB diluent solution (Dako, S3022) and incubated over night at 4 °C. Secondary ABs including DAPI and Phalloidin were incubated for 2 h/RT. After each AB incubation the sheet was rinsed 3× with PBS and washed 3× for 10 min. Finally the stained sheet was mounted in 50 μl Mowiol.

### Isolation of single Stratum Granular (SG) and Stratum Spinous (SS) cells

Dorsal back skin of newborn mice was excised (approx. 10mm x 10mm) and fat tissue was removed with fine curved forceps. The epidermis was mechanically, carefully peeled off from the dermis with ultrafine curved tweezers thereby separating the basal from the suprabasal layers. Subsequently, the suprabasal epidermis was floated on a 400 μL droplet of 44 μg/mL rETA (Staphylococcus aureus Exfoliative Toxin A)/1mM CaCl_2_ and incubated in the humidifying chamber at 37°C for 35 min. ETA diffuses intercellularly up to the tight junctions in the SG2 and cuts desmogleins-1 thereby loosening intercellular connections below the SG2 (Matsui et al., 2021). Layers below the SG2 layer (SS-SG3) were then peeled from the SG-1/2-corneum layers. Both parts were then floated onto 500 μL droplets of 0.05% trypsin/0.48 mM EDTA and incubated in the humidifying chamber at 37°C for 30 min. After the trypsinization, SS-SG3 cells were dissociated and SG-1/2 cells washed off from the stratum corneum by manual pipetting and transferred to a 1.5 mL Eppendorf tube. To neutralize, 1 mL of PBS++ was added to the trypsin solution containing SG and SS cells and centrifuged at 3000 rpm for 5 min at RT. For treatment, cells were resuspended in Dulbecco’s Modified Eagle Medium (DMEM) (Gibco).

### Cytoskeletal inhibitor treatment on isolated SG and SS cells

After isolation, cells were treated with 1 mL DMEM with or without Latrunculin B, 0.1 μM, shaking at 300 rpm for 1 hr at 37°C. After the treatment cells were washed with 1 mL of PBS++, centrifuged at 3000 rpm for 5 min at RT, and fixed with 4% PFA for 10 min at RT. Cells were permeabilized using 0.5% TritonX100/PBS for 10 min and washed with 500 μL of PBS++. Consecutively, cells were treated with Phalloidin-TRITC and Dapi for 30 min at RT. After the incubation, cells were centrifuged and mounted on coverslips with 30 μL of Mowiol. Cell were then imaged and the circumference of the cells was delineated using the freehand tool in Fiji. The area and circularity were extracted using the measure function in Fiji.

### Cytoskeletal inhibitor treatment on embryos

For *in vivo* Latrunculin and Y27632 treatments, wild-type embryos or *shScr; TRPV3-GFP* transduced embryos were collected on E17.5 with 2.5μM latrunculin (Sigma-Aldrich) or with 40μM Y27632 (Sigma-Aldrich) or DMSO in serum-free DMEM (Biological Industries) at 37⁰C for 1 h before embedding in OCT.

### Transglutaminase (Tgm) enzyme-substrate assay on cryosections

Cryosections were washed with PBS++ until the cryo tissue embedding compound Tissue-Tek O.C.T Compound (Sakura Finetek 4583) was removed. Then, sections were circled with a hydrophobic barrier marker pen (ReadyProbes™ - Thermo Fisher Scientific) and blocked with 10% NGS/PBS for 1 h at RT. Subsequently, sections were treated with and without Transglutaminase (Tgm) peptide substrate (pepK5: 5/6-FITC-Doa-YEQHKLPSSWPF-NH2/OH and K5pepQN: 5/6-FITC-Doa-YENHKLPSSWPF-NH2/OH, Intavis peptide services) for 30 min at RT in a humidified chamber and washed three times with PBS to remove the unspecific binding. Next, sections were fixed with 4% PFA for 5 min at RT and washed with PBS three times for 5 min. Section were mounted using mowiol.

### Toluidine blue barrier assay

Epidermal barrier assay was performed as previously described (Hardman et al., 1998). Briefly, E17.5 embryos were collected and immersed in an ice-cold methanol gradient in water, taking 2 min per step (1–25%, 2–50%, 3–75%, 4–100% methanol) and then rehydrated using the reverse procedure. Embryos were immersed in 0.2% toluidine blue solution. Embryos were washed in PBS before image capture.

### Microscopy

Confocal images were obtained with a Leica TCS SP8, equipped with gateable hybrid detectors (HyDs). Objectives used with this microscope: PlanApo 63x, 1.4 NA CS2. Images to be used for deconvolution were obtained at optimal resolution according to Nyquist. Alternatively, images were acquired with a Nikon C2+ laser-scanning confocal microscope using a 60x, 1.4NA oil or a 20x, 0.75NA air objective (Nikon). Epifluorescence images were obtained with a Leica DMI6000. Objectives used with this microscope: PlanApo 63x, 1.4 NA.

### Laser ablation

Keratinocytes were seeded on glass-bottom dishes (Ibidi # 81158) in FAD low Ca^2+^ medium. At confluency, the medium was switched to a high Ca^2+^ medium for 48 h. 2 h prior to imaging, medium was changed to high Ca^2+^ medium containing 1 μM SiR-actin (Spirochrome #CY-SC001). For laser ablation, a spinning Disk microscope (UltaView VoX, Perkin Elmer) equipped with a 355 nM pulsed YAG laser (Rapp Opto Electronic) was used. Experiments were performed at 37°C and 5 % CO2 using a water immersion objective 60 x, 1.2 NA (Nikon). The apical area apical stratified cells was ablated by drawing a line of consistent length with 30-50% laser power transmission and 10 iterations using UGA-40 Software (BioMedical Instruments).

### Transepithelial resistance measurement

A total of 5 × 10^5^ keratinocytes were seeded on transwell filters (Corning (#3460), 0.4 μm pore size). Cells were allowed to settle and then switched to 1.8mM high Ca2+ medium. Formation of TER was measured over time using an automated cell monitoring system (cellZscope, nanoAnalytics).

### Semiquantitative RT-PCR

RNA was extracted from samples using a Direct-zol RNA extraction kit (Zymo Research; R2060), and equal amounts of RNA were reverse transcribed using ProtoScript First Strand cDNA Synthesis Kit (New England Biolabs). Semiquantitative PCR was conducted using a StepOnePlus System (Thermo Fisher Scientific). Data are presented as mRNA levels of the gene of interest normalized to peptidylprolyl isomerase B (Ppib) mRNA levels. Primers used in this study: *Sptan1* Forward 5’-TCGACAAGGACAAGTCTGGC-3’; *Sptan1* Reverse 5’-AACAGGGCAAGCAGTGTAGG-3’; *Ank3* Forward 5’-CTGACGTTCACGAGGGAGTT-3’; *Ank3* Reverse 5’-TATCTAACGTGTCCGCTGCC −3’; *PPIB* Forward 5’-GTGAGCGCTTCCCAGATGAGA-3’; *PPIB* Reverse 5’-TGCCGGAGTCGACAATGATG-3’

### Image quantification

#### Quantification F-actin, αII-spectrin, pEGFR, Ankyrin3 and TGM1 substrate levels in embryos

Intensities from layers as indicated in the figures and legends was measured using the freehand selection tool (Image J) with a width of 5 pixels. All intensity measurements were determined by CTCF (Corrected Total Cell Florescence) = Integrated density-(Area X mean of fluorescence of background readings). Intensity levels in each layer were normalized according to controls.

#### Quantification of Cell shapes in embryos and newborns

Embryos that were injected with lentiviruses encoding *shScr;H2B-GFP,shsptan1;H2B-GFP* and *shAnk3-6780;H2B-GFP* on E8.5 and littermates were harvested at E17.5 frozen in OCT, sectioned (10 μm), fixed, and stained for E-cadherin (Cell Signaling Technology, 3195, 1:500). For the analysis of Ctr, *Cdh1^epi-/-^* and *Sptan1^epi-/-^* newborn epidermis, cryosections were PFA fixed and stained with a combination of Dsg1/2 antibody (Progen 61002, 1:200) and Dsg3 antibody (MBL D218-3, 1:2000). Samples were imaged using confocal microscopy. The cell boarders were delineated by the Freehand tool (Fiji) and cell area and perimeter using the “measure” function of Fiji. The cell shape index was calculated by dividing the perimeter by the square root of the area (Sahu et al., 2020b).

#### YAP+ cell quantification

The number of GFP+ and YAP+ cells was counted manually. The percentage of YAP+ cells was calculated as (number of YAP+GFP+ double-positive cells/total number of GFP+ cells) × 100 for basal and suprabasal layer.

#### Quantification of cortical enrichment

The mean gray value of the granular layer cells’ cortical and cytoplasmic intensity levels were measured with the “wand tool” (ImageJ) with a width of 5 pixels. The ratio between the two means was calculated and referred to as the fold change.

#### Statistics and repeatability of experiments

The numbers of independent experiments and biological replicates performed for all experiments, *p* values and the statistical tests that were used are indicated in the figure legends.

## Ethics declaration

All animal protocols in this study have been approved by either the animal experiment committee of LANUV, North Rhine-Westphalia, Germany or University Animal Care and Use Committee, confirmation number TAU-MD-IL-2206-162-4. All methods were carried out in accordance with relevant guidelines and regulations. The animal protocols and the reporting in this manuscript follow the recommendations in the ARRIVE guidelines.

## Data availability

The authors declare that the data supporting the findings of this study are available within the paper and its Supplementary Information files. Additional data are available from the corresponding author upon reasonable request.

## Author contributions

MR, CL, AS and CMN conceived of the study. MR, CL, AS, TM and CMN designed experiments. AS, AB, MP and MR carried out experiments and analyzed data. MR, CL, AS and CMN wrote the manuscript. All authors provided intellectual input, vetted and approved the final manuscript.

## Acknowledgements

We like to thank the CECAD imaging facility under supervision of Astrid Schauss. This work is supported by the Deutsche Forschungsgemeinschaft DFG, German Research Foundation: Germanýs Excellence Strategy – EXC 2030/CECAD – 390661388 (to CMN); CECAD project: CECAD Career-Promoting Grant for Senior Postdoctoral Scientists (to MR); SPP 1782 NI 1234/6-2 (to CMN); ISF grant number 145/23 (to CL). This work was carried out in partial fulfilment of the requirements for a Ph.D. degree for AS. from the Faculty of Medical & Health Sciences, Tel-Aviv University

